# Benchmarking imputation accuracy in the presence or absence of a reference panel

**DOI:** 10.1101/2025.10.08.680883

**Authors:** Alexandros Topaloudis, Tristan Cumer, Eleonore Lavanchy, Anna Hewett, Anne-Lyse Ducrest, Céline Simon, Bettina Almasi, Alexandre Roulin, Olivier Delaneau, Jérôme Goudet

## Abstract

Whole genome sequencing (WGS) of a large number of samples is costly. Solutions to reduce this cost include targeting a proportion of the genome (e.g. SNP arrays) or lowering the sequencing depth (low-coverage WGS, lcWGS) but both solutions suffer from either genotype missingness or uncertainty. Genomic imputation addresses this problem by inferring missing or uncertain genotypes using a collection of high quality genomic data (a reference panel). However, certain methods can impute a lcWGS dataset without a reference panel. Because the investment into generating a reference panel can be prohibitively expensive in non-model species, a benchmarking of the accuracy of these alternative methods of imputation can help inform study design. Here, we imputed a dataset of 2800 lcWGS in the presence and absence of a reference panel of 502 samples. We used 32 individuals sequenced at both high and low coverage to estimate the accuracy of each method and explored the limitations of lcWGS sample size and reference panel size. Although the best results were achieved with a large reference panel, using only a lcWGS dataset showed accurate imputation, when over 500 samples were used and we account for missing data and low frequency alleles. In addition, imputing with or without a reference panel gave similar results in a GWAS for a polygenic trait but required some quality control in the identification of homozygous-by-descent segments. Thus, while using a reference panel remains the ideal approach, imputation in suitably large lcWGS datasets can provide sufficient accuracy given proper quality control.

## Introduction

In the last couple of decades, the development of fast and affordable DNA sequencing techniques has enabled the rapid generation of genomic data which have become the backbone of many research fields, from medicine to evolutionary biology. Although the cost of DNA sequencing has reduced by multiple orders of magnitude, the errors that stem from sequencing technologies and downstream analyses (e.g. mapping, variant calling) usually necessitate sequencing a target sample in high coverage/sequencing depth (often at least 20x, Nielsen et al. 2011; Sims et al. 2014). As a consequence, sequencing can still come at a daunting cost when the number of samples required is large. Solutions to reduce these costs include sequencing a fraction of the genome (e.g. SNP arrays, Restriction site Associated DNA sequencing (RADseq)) or sequencing the whole genome at a fraction of the depth (e.g. 1 - 2x) an approach termed low-coverage whole genome sequencing (lcWGS) (Fuentes-Pardo and Ruzzante 2017; Benjelloun et al. 2019; Lou et al. 2021). SNP arrays have been widely used for model species but they bear a large up-front cost to develop, require a reference genome and a catalog of existing variation, and are not readily transferable across populations (Nielsen 2004; Ha et al. 2014). RADseq offers an alternative for non-model species but also comes with pitfalls such as a biased sampling along the genome (Davey and Blaxter 2010) and a lower repeatability (Cumer et al. 2021) which affect downstream analyses (Lavanchy and Goudet 2023). Similarly, lcWGS has its own limitations: it requires an assembled reference genome (although more and more reference genomes for different species are becoming available; (Rhie et al. 2021; Gupta 2022; The Darwin Tree of Life Project Consortium 2022)) and comes with significant genotypic uncertainty and high levels of missing data. In contrast,, lcWGS enables the de-novo discovery of variation in a sample of individuals (Le and Durbin 2011; Martin et al. 2021) and there exist specialised methods and tools that can account for the substantial genotypic uncertainty (e.g. ANGSD, Korneliussen et al. 2014) making lcWGS an attractive solution for non-model species (Lou et al. 2021).

To overcome the shortcomings of both SNP arrays and lcWGS, the data produced can be coupled with genomic imputation, which generates genome-wide data (often phased) from incomplete observations, providing access to a powerful set of analyses (Marchini and Howie 2010; Das et al. 2018). Usually, genomic imputation infers unobserved genotypes, such as variants untyped in SNPs array data and uncertain or missing in lcWGS by using a reference set of high quality haplotypes (a reference panel). The principles behind imputation lie in population genetic theory (Li and Stephens 2003; Browning 2008; Marchini and Howie 2010; Das et al. 2018) and utilise the property that even distantly related individuals in a population share segments along their genome that are identical-by-descent (IBD). Therefore, the unobserved haplotypes of a focal sample can be ‘pieced together’ by using the scarce genotyped data available and identifying the corresponding observed haplotypes present in the reference panel. The algorithms account for differences due to mutation and allow haplotype switching due to recombination events and are usually modelled under a hidden Markov model. Different versions of imputation have been implemented using different input data and assumptions (e.g. BEAGLE, Browning et al. 2018; GLIMPSE, Rubinacci et al. 2020; QUILT, Davies et al. 2021; IMPUTE5, Rubinacci et al. 2021). Because they rely on linkage disequilibrium (LD), these methods can be collectively classified under the term ‘linkage-disequilibrium imputation methods’. In addition to methods mentioned above, some implementations can take advantage of specialised resources available in the target species. For example in livestock species where large pedigrees are available specialised imputation software can utilise relatedness along a pedigree to impute target individuals (e.g. (Hickey et al. 2012; Cheung et al. 2013; Sargolzaei et al. 2014).

Genomic imputation has proven to be remarkably accurate. Through testing using empirical and simulated datasets, the inferred imputation accuracy is very high for inferences in human and livestock genetics as well as in ancient DNA (Pasaniuc et al. 2012; Lloret-Villas et al. 2023; Sousa da Mota et al. 2023; Erven et al. 2024). Specifically, imputation performance tends to be excellent for alleles at intermediate frequencies but diminishes for rare alleles, although including pedigree information can improve imputation of low frequency alleles (Liu et al. 2019). The negative correlation of imputation accuracy and allele frequency stems from rare alleles being found in only few copies in the reference panel haplotypes and not being sufficiently tagged by surrounding variation making them harder to infer. In addition to allele frequencies, imputation performance is sensitive to the composition of the reference panel used. It has been shown that imputation performs best when the reference panel used is large, well-phased and is representative of the haplotypic diversity in the target population (Das et al. 2018; Garcia et al. 2022; Marino et al. 2022). These findings have spurred the development of vast reference panels in humans and other model species, where whole genome sequences of hundreds of thousands of samples are routinely used to impute from a scarcer set of typed variation (McCarthy et al. 2016; Rowan et al. 2019; Wang et al. 2022; Shi et al. 2024).

When a reference panel is not available, imputation can still be performed using only low-coverage whole genome sequencing data. The methods that accomplish this work in similar ways to the LD-based imputation methods but make assumptions about the underlying haplotypes that have to be pieced together by the lcWGS dataset instead of being observed in a reference panel. Existing implementations include STITCH (Davies et al. 2016), earlier versions of BEAGLE (Browning and Browning 2007) and SHAPEIT (Delaneau and Marchini 2014) among others, and have been called ‘genotype refinement’ in order to tell them apart from imputation methods using a reference panel (Das et al. 2018). The resulting accuracy of genotype refinement is promising and can pave the way for the wider adoption of lcWGS in wild populations, extending the possibilities for identifying genomic regions under selection, runs of homozygosity and other genome-wide patterns or for dissecting the genetic architecture of traits using Genome Wide Association Studies (GWAS), without the need of a costly reference panel (Delaneau and Marchini 2014; Davies et al. 2016; Teng et al. 2022; Yang et al. 2024). Nevertheless, the adoption of imputation in non-model species is only in its infancy, with few empirical studies using imputation of lcWGS in wild populations with or without reference panels (Fuller et al. 2020; Gao et al. 2021; Enbody et al. 2023; Hooper et al. 2024; Justen et al. 2024; Landis et al. 2024). This can stem both from the lack of available genomic resources but also from a lack of thorough validation of imputation methods in wild populations, which might generate uncertainty about the accuracy of the resulting genotypes. To date, there are very few tests of the accuracy of imputation in wild populations (Watowich et al. 2023; Tan et al. 2025), with none specifically comparing the presence and absence of a reference panel. While similar comparisons exist for humans and livestock (Shi et al. 2019; Teng et al. 2022; Wang et al. 2024), the low effective population size, high LD and unique mating schemes practiced in selective breeding can inflate imputation accuracy and these results are not readily transferable to wild populations. Furthermore, most benchmarking studies rely on simulated data as the ground truth or on artificially downsampling high coverage samples to generate in-silico a lcWGS dataset which might not accurately reflect the specificities of DNA sequencing.

To address this, we quantify the accuracy of imputation in a newly generated dataset of lcWGS in a wild population of barn owls (*Tyto alba*) where a reference panel is also available (Figure 1). We use 2800 samples recently sequenced in lcWGS (average depth = 2x - Figure 1B)(Cumer et al. 2025) and impute them with or without the use of a reference panel of 502 medium to high-coverage (hcWGS) individual whole genomes (Figure 1A) using GLIMPSE and STITCH respectively. We measure the imputation accuracy using 21 individuals sequenced both at high and low coverage (Figure 1C). We compare the true genotypes of these 21 individuals to the imputed genotypes obtained using a reference panel through GLIMPSE (2800 lcWGS + 502 hcWGS) and an imputed dataset through genotype refinement obtained using only the 2800 lcWGS in STITCH. Further, we evaluate the effect of reference panel size, and lcWGS sample size on imputation accuracy; and estimate the downstream impact on estimating inbreeding along the genome and carrying out a GWAS for a polygenic trait (Figure 1D). We explore the benefits and limitations of the two methods of imputation and provide guidelines on using lcWGS coupled with imputation.

**Figure 1.**
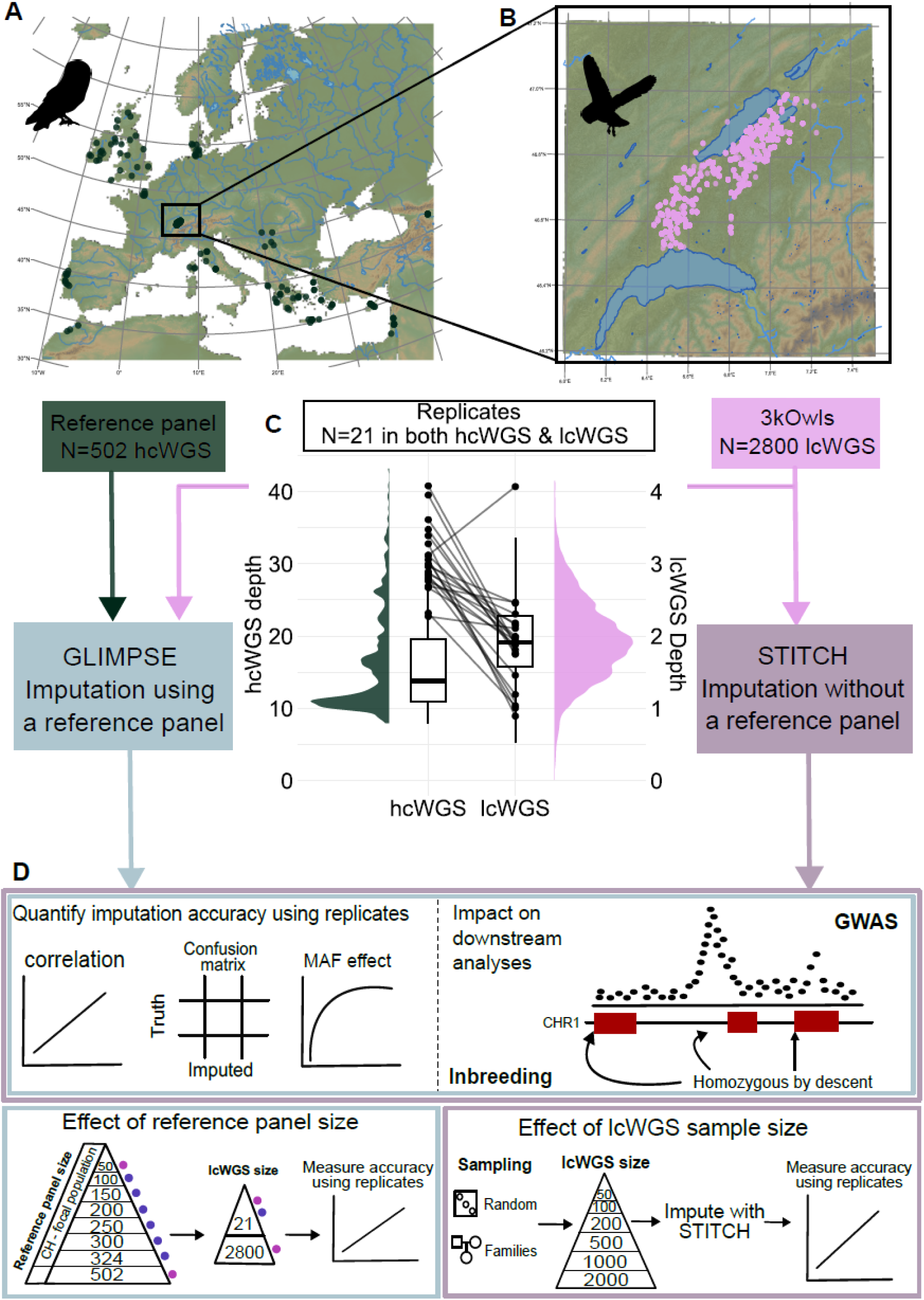
A graphical abstract of the study. **A)** The barn owl (*Tyto alba*) data consists of a reference panel of 502 samples sequenced in medium to high-coverage WGS (hcWGS), 346 of which come from the focal long term study in Western Switzerland (CH). **B)** 2800 samples from the long term study in Western Switzerland sequenced at low-coverage WGS (lcWGS). **C)** Sequencing depth of the datasets used. Left: sequencing depth in the reference panel; right: sequencing depth of the lcWGS. Black dots represent the 21 replicate samples sequenced in both datasets. **D) Top left**: the accuracy of imputation with GLIMPSE and STITCH quantified using the replicate samples including the effect of minor allele frequency (MAF). **Top right**: sketch of downstream analyses like a polygenic trait GWAS and inbreeding along the genome. **Bottom left**: The effect of reducing the reference panel size. Colored dots show which reference panel size was used to impute which lcWGS size (21 or 2800). **Bottom right**: The effect of lcWGS sample size and sampling strategy.

## Methods

### Reference panel

The set of individuals used as a reference panel in this study were initially sequenced in multiple past studies (Cumer, Machado, Dumont, et al. 2022; Cumer, Machado, Siverio, et al. 2022; Machado, Cumer, et al. 2022; Machado, Topaloudis, et al. 2022; Cumer et al. 2024; Topaloudis et al. 2025). Whole genome sequences of 502 individuals were genotyped together through a variant discovery pipeline briefly described below (full details can be found in the original publication where the dataset was generated, Topaloudis et al. 2025). Raw reads were processed with trimmomatic v0.39 (Bolger et al. 2014) which removed sequencing adapters and excluded unpaired reads and reads with a length smaller than 70bp. Mapping was performed with BWA-MEM v0.7.17 (Li and Durbin 2009) on the barn owl genome assembly v4 (Machado, Cumer, et al. 2022). Variant discovery followed the GATK v4.2.6 (Auwera et al., 2013) pipeline with a base quality score recalibration (BQSR) step performed using a previously published “truth set” of variation (Cumer, Machado, Dumont, et al. 2022). GATK’s *HaplotypeCaller* ran with default parameters for each individual followed by a common joint variant calling using *GenotypeGVCFs*. SNPs were then filtered to be bi-allelic, pass technical filters (QD<2.0, QUAL<30, SOR>3.0, FS>60.0, MQ<40.0, MQRankSum<-12.5 and ReadPosRankSum<-8.0) and further filtered to exclude regions of the genome where the mappability of the reads is limited (see Corval et al. 2023 for details). Further, genotypes were filtered on individual depth using *BCFtools* v.1.15.1 (Danecek et al. 2011). Any genotype call with a read depth lower than 5 or higher than three standard deviations above the individual average depth was set to missing. After these filters, the reference panel consisted of 10,451,268 SNPs. For the purpose of this study we only retained variants present on the 39 autosomal linkage groups of the barn owl genome (Topaloudis et al. 2025), resulting in a total of 10,115,035 autosomal SNPs.

Reference panel pre-phasing is required by most recent software so we phased the reference panel in two steps. First, individual variants were phased using an observational approach in which reads or pairs of reads covering multiple heterozygous sites were used to resolve local haplotypes (read-based phasing). Since some individuals come from families in the pedigree, the read information was used alongside family trio information (genetic phasing of a couple and its offspring) when available. For both read-based and trio-based (genetic) phasing we used *WhatsHap* v1.4 (Garg et al. 2016; Martin et al. 2016) with the mean recombination rate estimated for the species (--*recombrate* 2). All individuals belonging to one (or more) trio were phased using *WhatsHap’s pedigree-mode*. Offspring were phased with their parents, and parents which had more than one offspring were phased along with the highest sequencing depth offspring. After read-based and genetic phasing, we performed an additional step of filtering. We identified 187 unrelated individuals (kinship < 0.03125) and retained bi-allelic SNPs with minor allele count larger than five (MAC > 5) and missing data < 5% based on this set of unrelated individuals to account for non-independence of sampling in the family data.

This set of filtered variants was then statistically phased with SHAPEIT v4.1.2 (Delaneau et al. 2019). We phased the 187 unrelated individuals and 315 individuals belonging to families separately to avoid effects of family structure in the dataset. First, the 187 unrelated individuals were phased together and then each family member was independently phased with the unrelated set. We ran SHAPEIT following the authors’ instructions for increased accuracy: the number of conditioning neighbors in the PBWT was set to 8, and the MCMC chain ran with 10 burn-in generations, 5 pruning iterations, each separated by 1 burn-in iteration, and 10 main iterations.

The quality of the phasing was assessed using the switch error-rate (SER) (Browning and Browning 2011) metric: when comparing two phase sets for an individual’s variants, a switch-error occurs when a heterozygous site has its phase switched between the phase sets obtained with different phasing algorithms. For each individual, the local phasing inferred from WhatsHap (read-based approach and when possible genetic approach) was considered as the ground truth. This phase set was compared with the statistical phasing of this individual using Shapeit, with read-based/genetic phase information ignored for the individual. The final estimation of the switch error rate was done using the switchError code to compare both phasing sets (available at https://github.com/SPG-group/switchError). The results showed that the phasing of variants present in the reference panel yielded a low error rate (mean error rate of 1.83%, (Cumer et al. 2025))

### Low coverage dataset

We sequenced 2,800 owls from a pedigreed population in Switzerland (Roulin et al. 1998; Frey et al. 2011). Blood samples were collected between 1986 and 2020 and individuals with complete family information and phenotypic data available were prioritised for sequencing. For verification of sequencing performance, we included 32 individuals that had already been sequenced at high coverage in the reference panel. These individuals sequenced at both high and low coverage were used to measure the accuracy of imputation strategies.

For the lcWGS, genomic DNA was extracted from blood samples using the DNeasy Tissue Kit (Qiagen, Switzerland) and the Biosprint robot 96 (Qiagen, Switzerland) and stored at −20°C. We randomized the position of the DNA samples of the 2,820 owls on 30 different 96-well plates. Every plate included two empty wells to measure potential contamination. The DNA was quantified using Quant-it PicoGreen dsDNA Assay kit (Thermo Scientific, Switzerland) and was diluted in 10 mM Tris-HCl to 1.5 to 2.5ng/μl. The libraries were prepared with 8 plexWell 384 kits (SeqWell, USA) and sequenced in 3 flow cells of an Illumina NovaSeq 6000 at the genomic technologies facility of the University of Lausanne (GTF).

Raw reads were trimmed with *Trimmomatic* v.0.36 (Bolger et al. 2014) and aligned to the barn owl reference genome v4 (Machado, Cumer, et al. 2022) using *BWA-MEM* v.0.7.15 (Li and Durbin 2009). Mean coverage of the genome was 1.95X with a minimum coverage of 0.2X and maximum coverage of 4.15X (Figure 1C).

### Imputation

#### GLIMPSE - imputation using a reference panel

The imputation using GLIMPSE is described in detail in Cumer et al., (2025).

We determined genotype likelihood at each variant position present in the reference panel using *BCFtools* v.1.15.1 (Danecek et al. 2021) *mpileup* and *call* methods, using the -T and -C options. We then used *GLIMPSE* v1.1.1 (Rubinacci et al. 2021) for the imputation and phasing of the low coverage individuals using the reference panel described above. We used the first version of GLIMPSE (v1.1.1) because it is more suitable for small reference panels according to the GLIMPSE manual. The *GLIMPSE* pipeline was applied in four steps: The first step, called chunking (implemented in the *GLIMPSE_chunk* method), splits the chromosome into chunks for efficient imputation and phasing. We performed this chunking with default parameters (*--window-size* 2000000 and *--buffer-size* 200000). The second step consists of the phasing and imputation of the genomic chunks (implemented in the *GLIMPSE_phase* method). This method iteratively improves low-coverage genotypes’ likelihoods and phasing for each individual independently. We conducted phasing and imputation with the *GLIMPSE_phase* method, with the maximum number of iterations (--*burnin* 100 and --*main* 15) and with a large effective population size parameter value (--*ne* 10000). The recombination map was taken into account with the --*map* parameter. The third step is the ligation step which merges the different chunks without losing phase information. This procedure is implemented in the *GLIMPSE_ligate* method which was executed with default parameters. The last step of the *GLIMPSE* pipeline is haplotype identification (implemented in the *GLIMPE_sample* method) which identifies the most likely haplotype based on posterior genotypes’ likelihood and phase information (we run this using the --*solve* mode).

Because including an individual both in the reference panel and the imputation target would bias the results we ran the *GLIMPSE* pipeline three times: once with the full reference panel (the 502 individuals) and with the 2,768 unique low-coverage samples. We then removed the 22 duplicated samples from the reference panel and included them in the low-coverage set and ran the *GLIMPSE* pipeline again (with a reference panel of 480 individuals and a low coverage data set of 2,790 individuals). We also ran *GLIMPSE* separately for the 10 other replicates (with a reference panel of 492 individuals and a low coverage data set of 2,778 individuals). Lastly, we merged the core dataset of 2,768 individuals with the 22 and 10 replicates, into a total dataset of 2,800 unique imputed samples.

For subsets of the reference panel, the 22 replicates were imputed with a random selection of 50,100,150,200,250,300 samples from Switzerland, as well as with the whole Swiss dataset (346 in total - 22 replicates = 324 samples) and with the full reference panel (502 - 22 = 480 samples). All other parameters were kept the same as in the full run.

#### STITCH - imputation without a reference panel

STITCH v1.7.2 (Davies et al. 2016) was used to impute the genotypes of the 2,800 low coverage samples. The software models the population being founded by *K* founders *n* generations ago, and requires a list of positions to genotype in all samples. In all runs *n* can be defined as 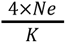, with *Ne* being the effective population size (in our case we used 10,000 in line with the GLIMPSE implementation). The optimal number of *K* is case-dependent and needs to be determined. We chose two linkage groups with strikingly different recombination landscapes, one with a punctuated landscape of recombination (LG22, 165k SNPs, 25Mb in length, ∼2.5 cM/Mb) and one with elevated recombination across its length (LG36; 80k SNPs, 7.8Mb in length, ∼6.4 cM/Mb) (Topaloudis et al. 2025). We genotyped SNP positions present in the reference panel and ran STITCH using the ‘diploid’ method on the full dataset of 2,800 samples for values of K ranging from 4 to 40 with a step of 2. We then compared the squared correlation of the inferred genotypes with the ‘true’ genotypes for the 32 replicate individuals to identify an optimal value of K (Supplementary Figure 1). We chose K=30 as a tradeoff of running time and accuracy which we used for all downstream runs of STITCH.

To run STITCH a set of variant positions are needed. In order to follow a similar pipeline as a new study without a reference panel would, we decided not to use the variants present in the reference panel but to perform snp-calling de-novo. To this end we performed variant calling in the low-coverage samples using *BCFtools* v.1.15.1. The genotypes were generated with the *bcftools mpileup* command with the following parameters (-B, -q 20 -Q 20 -a ‘INFO/AD,AD,DP’) using the barn owl reference genome v4 and the mappability mask mentioned above (see Reference Panel). Then variant positions were called using the *bcftools call* command with calling indels (-V indels) on a multivariate caller mode (-mv). Variants were filtered for missingness (maximum 30%), genotype quality (minimum 20) and total depth (minimum 1000) and made into a position file for use in STITCH using *bcftools query*. Then STITCH was run in 5Mb windows setting K to 30, and the number of generations to 1350 with the method set to the diploid model for 40 iterations and all other parameters as default.

To test for the effect of sample size we subsampled the low coverage datasets to sizes of 50, 100, 200, 500, 1000 and 2000. We always included the 32 replicated individuals and the rest of the individuals were picked by either random sampling or by sampling individuals prioritising families. Families were defined by sharing a family ID generated with the *makefamid* function in the R package *kinship2* v1.9.6 (Sinnwell et al. 2014). For each subset we always chose the largest possible family to include, and repeated the process until the subset was filled.

We also imputed the lcWGS dataset without a reference panel using a combination of ANGSD, for SNP calling (Korneliussen et al. 2014) and BEAGLE, for imputation (Browning and Browning 2009) as described in the ANGSD manual. Because STITCH outperformed ANGSD/BEAGLE, we only present the results for STITCH in the main text but the results for BEAGLE can be found in the Supplementary Material.

### Downstream comparison

#### Statistics

Using the 32 individuals present in both high and low coverage we applied different metrics to quantify the performance of imputation. The accuracy of imputation, r^2^, is the squared correlation of the genotype vectors for each individual between the true (high coverage) and the imputed dataset. In addition we estimated the imputation accuracy on the centered and scaled genotype vectors as suggested in Calus et al. (2014). The scaled imputation accuracy gives larger weight to rare alleles and results were identical in rank but smaller in absolute value when using the scaled correlation (Supplementary Figure 2). In all comparisons, the reference allele was chosen to be the most common across all methods. Sensitivity (true positive rate) refers to the ratio of variants in a genotype class correctly inferred to be in that genotype class (i.e. the ratio of true positives to the sum of true positives and false negatives). Precision (positive predictive value) is the flipside of sensitivity as it measures the ratio of variants inferred to be in a genotype class truly being in that genotype class (i.e. the ratio of true positives to the sum of true positives and false positives). All statistics were estimated from comparing the genotype matrices generated either from GLIMPSE or STITCH or by extracting the information from the *vcf* files. Since GLIMPSE does not report missing genotypes we simply converted the *vcf* file into dosage format. For genotypes set to missing in STITCH, we extracted the genotype dosage from the *GT:DOS vcf* field as reported by STITCH. For STITCH, float genotype values from likelihood calls were rounded to the nearest integer for comparison with the integer high coverage genotypes and all statistics were estimated using the *confusionMatrix* function of the R package *caret* v.6.0-94 (Kuhn and Max 2008). All imputation software output an accuracy metric per variant reflecting the confidence in the imputed genotypes. The metric used in GLIMPSE and STITCH is the imputation ‘information score’ (info score). All individual accuracy measures were estimated for three INFO cutoff filters (0, 0.6, 0.8) and with or without taking into account the missing genotypes in STITCH.

To identify the source of variation in per sample imputation accuracy we regressed the per sample imputation accuracy of each high-coverage replicate (n=21) to the number of close relatives (relatedness > 0.1) in the reference panel or the lcWGS dataset and the sequencing depth in high or low coverage, using the *lm* function in R.4.4.1. Relatedness was calculated using a relatedness matrix of the samples estimated with *hierfstat* v.0.5-11 (Goudet and Jombart 2022).

To quantify per variant imputation accuracy we compared the genotype vectors of the 21 replicates sequenced in the highest coverage (> 20x). To quantify imputation accuracy in rare alleles we used the non-reference-concordance (NRC). We define NRC as 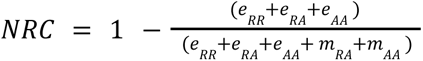, where *e_x_* refers to the number of mismatches of genotypic class ‘x’ and *m_x_* refers to the number of matches in genotypic class ‘x’. Genotypic classes are defined as RR: homozygous for the common allele, RA: heterozygotes, AA: homozygous for the rare allele. We quantified NRC for each set of variants in the following allele frequency bins “0.00001 0.00100 0.00200 0.0030 0.0040 0.00500 0.01000 0.01500 0.02000 0.02500 0.03000 0.04000 0.05000 0.10000 0.20000 0.30000 0.40000 0.50000” using the *GLIMPSE_concordance* tool of GLIMPSE v1.1.1. The tool was run using the genotype mode *(--gt-validation,--gt-target*) for each imputed dataset. We quantified allele frequencies in 4 datasets: 76 unrelated Swiss samples present in the reference panel, the whole reference panel (n=502), the imputed dataset from GLIMPSE (n=2800) and the imputed dataset from STITCH (n=2800). Allele frequencies were highly correlated and most accurate in large sample sizes as expected and for the comparison of NRC per MAF bin we used the allele frequencies estimated in GLIMPSE (Supplementary Figure 3).

#### Homozygosity-by-descent estimation

To generate estimates of homozygosity-by-descent (HBD) and annotate the origin of HBD segments along the genome we used the RzooRoH v0.4.1 R package (Bertrand et al. 2019). A subset of 51 individuals (21 replicates and 30 individuals previously shown to be inbred (Hewett et al. 2025) was used. The 30 inbred individuals sequenced on lcWGS were included to quantify the relative performance of imputation in more highly inbred individuals. We firstly filtered for INFO > 0.8 and an allele frequency of 0.1%, then ran the default RzooRoH model with the number of K-rates set to 14, corresponding to HBD origins of 2, 4, 8, 16, 32, 64, 128, 256, 512, 1024, 2048, 4096 and 8192 generations ago. For the high-coverage dataset, we ran RzooRoH using the same parameters but including only the 21 replicates.

#### GWAS

To test the performance of the imputed dataset in a genome wide association study we used the mean tarsus length (mm) of adult birds, a polygenic phenotype according to preliminary analyses, with intermediate heritability in adults (Hewett et al. 2025). A total of 1980 individuals were included in the GWAS. A linear mixed model was used in the *gaston* R package v.1.5.9 (Perdry et al. 2018) with the function *association.test*. Sex was fitted as a fixed effect, formatted as a factor. In addition an allele-sharing (Weir and Goudet 2017; Goudet et al. 2018) kinship matrix estimated with *hierfstat* v.0.5-11 (Goudet and Jombart 2022), doubled to correspond to a relatedness matrix, was used as a random effect to control for structure and relatedness in the dataset.

All R-related analyses were executed under R version 4.4.1. All scripts for the analyses can be found in the following GitHub repository: https://github.com/topalw/Imputation_owls.

## Results

### Comparison of approaches

To test the accuracy of imputation in a wild population with and without a reference panel we imputed 2800 low-coverage whole genome sequences (lcWGS) with and without the use of 502 medium to high coverage WGS (hcWGS) (Figure 1). When making use of the hcWGS, we used the software GLIMPSE, which imputed a set of 10,102,233 bi-allelic SNPs previously identified in the hcWGS and genotyped in the lcWGS data. When using only the lcWGS dataset, we used STITCH which imputed a set of 24,224,697 bi-alllelic SNPs de-novo identified in the 2800 lcWGS samples.

For validation we used a set of 32 samples sequenced both at high and low coverage, which we call ‘replicates’. All high coverage genotypes of the replicates were removed from the reference panel during imputation with GLIMPSE (see Methods). Of the 32 samples, one was a library error and is excluded from all analyses. The remaining 31 samples showed variable performance in imputation accuracy. Specifically 10 samples sequenced at lower sequencing depth in the high coverage ‘truth set’ (mean depth 11x; range 8.4 - 17.3x; hereafter ‘intermediate depth’) showed a pattern of very high sensitivity (true positives over true positives and false negatives) and low precision (true positives over true positives and false positives) only in the heterozygote genotypes (Supplementary Figure 4 & 5). This pattern could stem from missing heterozygous calls in the ‘truth dataset’ which were then correctly inferred when imputation was used but classified as errors in the comparison. Consistent with this hypothesis, the 10 samples in intermediate depth exhibited less heterozygous sites than the 21 higher depth samples (Supplementary Figure 6). For this reason these 10 individuals were subsequently excluded from the replicate set used to validate the imputation results. Thus, unless stated otherwise, we use 21 samples whose high coverage genotypes (mean depth = 30.5, range = 22.7 - 40.8, Figure 1C) are considered the ground truth against which all comparisons are made, unless stated otherwise.

We tested the effect of filtering on the accuracy of imputation by using only SNPs with an information score larger than 0.6 or 0.8 compared to using the unfiltered set of variants. Both cutoffs increased the quality of imputation at the cost of reducing the number of variants remaining (Supplementary Figure 7). Filtering at 0.8 reduced the number of variants retained in GLIMPSE and STITCH to 94% and 83% of the method’s total, respectively (Supplementary Table 1). For the rest of the results we use an imputation accuracy filter of 0.8 leaving 7,182,235 SNPs at the overlap of GLIMPSE and STITCH. Results for different filtering cutoffs can be found in Supplementary Figures 8 & 9.

### Overall imputation accuracy

The per sample accuracy of imputation (quantified as the squared Pearson correlation, r^2^, between the high coverage and imputed genotypes of each replicate) was very high (> 0.95) regardless of whether a reference panel was used or not (Figure 2A). However, using a reference panel with GLIMPSE showed significantly higher imputation accuracy (mean=0.978; sd=0.008) with lower variation among individuals, compared to STITCH (mean=0.966; sd=0.014; Wilcoxon rank sum exact test; *W* = 339, *p* = 0.002).

**Figure 2.**
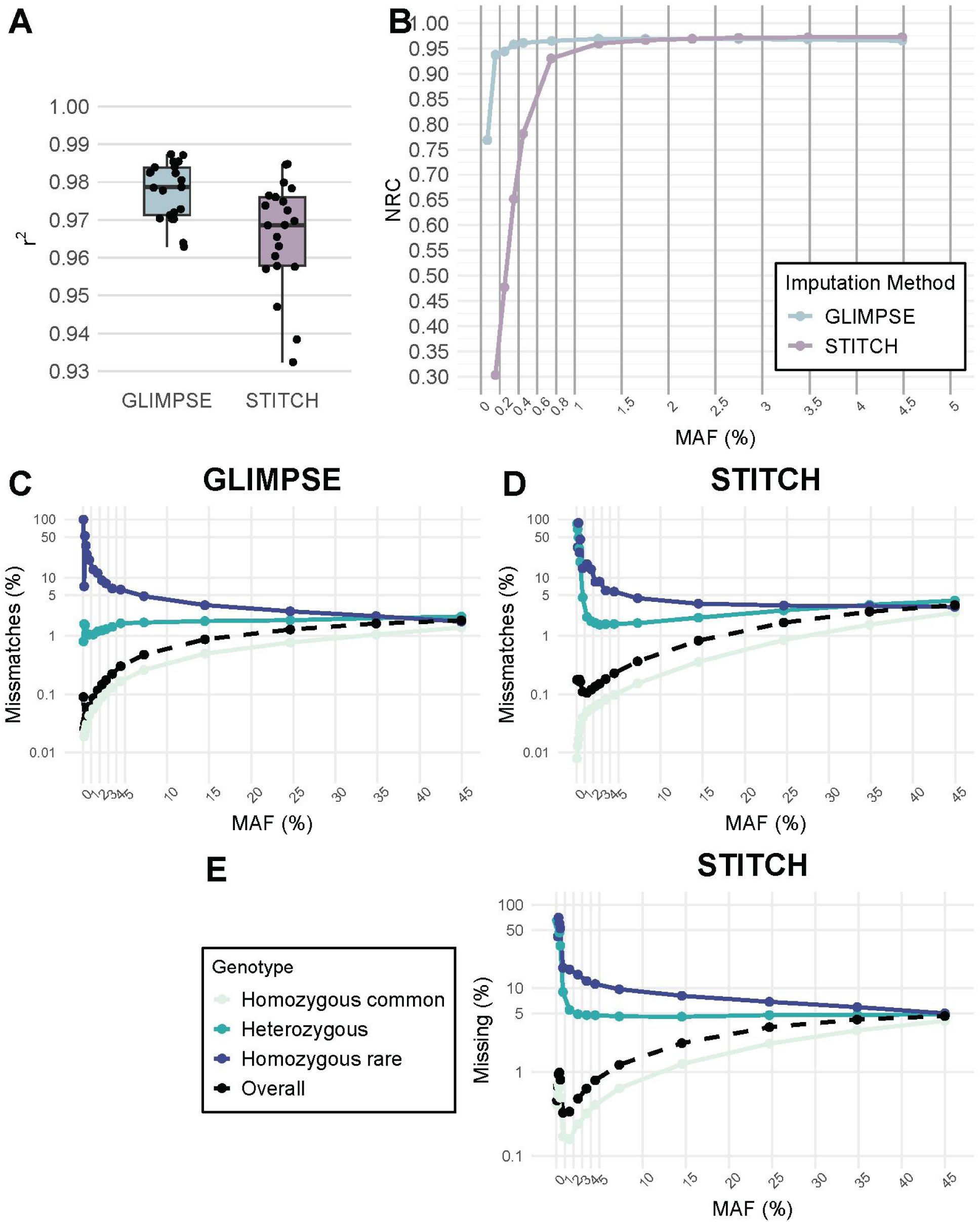
The accuracy of imputation with and without a reference panel. A) the squared correlation (imputation accuracy) per sample between the ‘truth’ (high-coverage data) and the low-coverage WGS dataset imputed by GLIMPSE or STITCH. Each point is one of the 21 replicate individuals. B) Non-reference concordance (NRC) per minor allele frequency bin. Each point represents the mean in the specific allele frequency bin. Allele frequency for the rare allele was defined in the GLIMPSE dataset of 2800 samples. C-D) The misclassification profile of each genotype class across each allele frequency bin for GLIMPSE C) and STITCH D). E) Missing data profile for STITCH for each genotype class across each allele frequency bin. Legend shared across panels C,D and E.

We then investigated what could explain the among-sample variation in imputation accuracy. We regressed the GLIMPSE imputation accuracy of the 21 replicates on the sequencing depth in the truth and the low-coverage set along with the number of close relatives (relatedness > 0.1) in the reference panel and the lcWGS dataset. The model explained 8.5% (adjusted R^2^ = 0.085) of the variation in imputation accuracy and no factor was significant. For STITCH we repeated the same model excluding the number of relatives in the reference panel. This model explained approximately 25% of the per-sample variation in imputation accuracy (adjusted R^2^ = 0.246). The only statistically significant factor in the model was the low-coverage sequencing depth of the sample (β=0.02, p<0.01). In both models, including the 10 replicates sequenced in intermediate depth, showed a strong effect of the high-coverage sequencing depth, in line with the assumption that lower depth samples underestimate imputation accuracy due to errors in the truth set (Supplementary Figure 10-12).

Imputing rare alleles is notoriously difficult (Marchini and Howie 2010). To measure performance across allele frequencies, we quantified the non-reference concordance (NRC) which is better suited for low-frequency alleles than imputation accuracy where the multiple homozygotes for the common allele will bias the results (Li et al. 2011; Bizon et al. 2014). When referring to a reference allele we always refer to the most common allele at each locus. Both STITCH and GLIMPSE showed very high NRC in allele frequencies above 1-2% (Figure 2B). When using STITCH, NRC dropped slightly for variants with a minor allele frequency below 1.5% and plummeted below 0.8%. The use of a reference panel with GLIMPSE showed accurate estimation across most allele frequency classes but exhibited a sharp decrease in allele frequencies below 0.2%.

To identify the origin of imputation errors, we measured the proportion of each genotype misclassified with each method along the allele frequency bins (Figure 2C, 2D). Both methods had a similar misclassification profile for all genotype classes, with STITCH exhibiting an increased overall rate across all frequency bins and genotype classes along with a higher misclassification of heterozygote genotypes in rare alleles (<1%). As expected, heterozygotes were misclassified as either homozygote genotypes and homozygotes for the alternative allele are almost always misclassified as heterozygotes (Supplementary Figure 13).

When using STITCH some genotypes are set to missing when the inferred genotype probability lies below a certain threshold (0.9 by default). This behaviour leads to a proportion of missing data per sample (average across 2800 individuals: 2.7%, range: 1.6% - 25%) which correlates strongly and negatively with the lcWGS depth each sample is sequenced in (Supplementary Figure 14). In addition overall missingness increased as the allele frequency increased (Figure 2E). For heterozygotes, high missingness was observed mostly in rare alleles and it plateaus to about 5% in allele frequencies larger than 2%. Lastly, the genotype class with the largest proportion of missing data is the homozygotes for the rare allele. However, because STITCH reports the genotype probability, even when the called genotype is missing, we could compare the full genotype probabilities (no missing data) with the true genotype of the replicates. The accuracy of imputation with STITCH per sample when using the genotype probabilities (including the non-called genotypes) dropped on average by 0.13% (using genotype calls, mean = 96.6%; using genotype probabilities, mean = 96.46%) (Supplementary Figure 15).

### The effect of reference panel size

To address the effect of reference panel size, we quantified imputation accuracy in subsets of the full reference panel. We ran GLIMPSE using a set of 324, 300, 250, 200, 150, 100 and 50 samples from the focal population of Western Switzerland. Using the above reference panel sample sizes, we imputed only the 21 replicate individuals. We also imputed the 21 replicates using the full reference panel of 502 individuals (480 if the 22 duplicates are excluded) but without other lcWGS samples. We observed increasing imputation accuracy with increasing reference panel size (Figure 3A). All reference panel sizes showed an average imputation accuracy above 0.9, with a reference pane size above 200 samples returning more than 0.95 imputation accuracy per sample. The largest benefit was seen when increasing the reference panel size from 50 to 200 individuals (1.15% accuracy increase per 50 samples added) while further increase provided reduced benefits (0.23% accuracy increase per 50 samples). Including the 156 non-Swiss samples in addition to the 324 Swiss led to a 0.6% increase in imputation accuracy. Imputing with the full reference panel using the 2800 lcWGS dataset, instead of only the 21 samples, led to a further 1.5% increase in imputation accuracy. Non-reference concordance increased with increasing reference panel size across all allele frequencies (Figure 3B), but in particular for allele frequencies below 0.5% and the inclusion of a larger number of lcWGS produced better results for allele frequencies between 0.1 and 0.5%.

**Figure 3.**
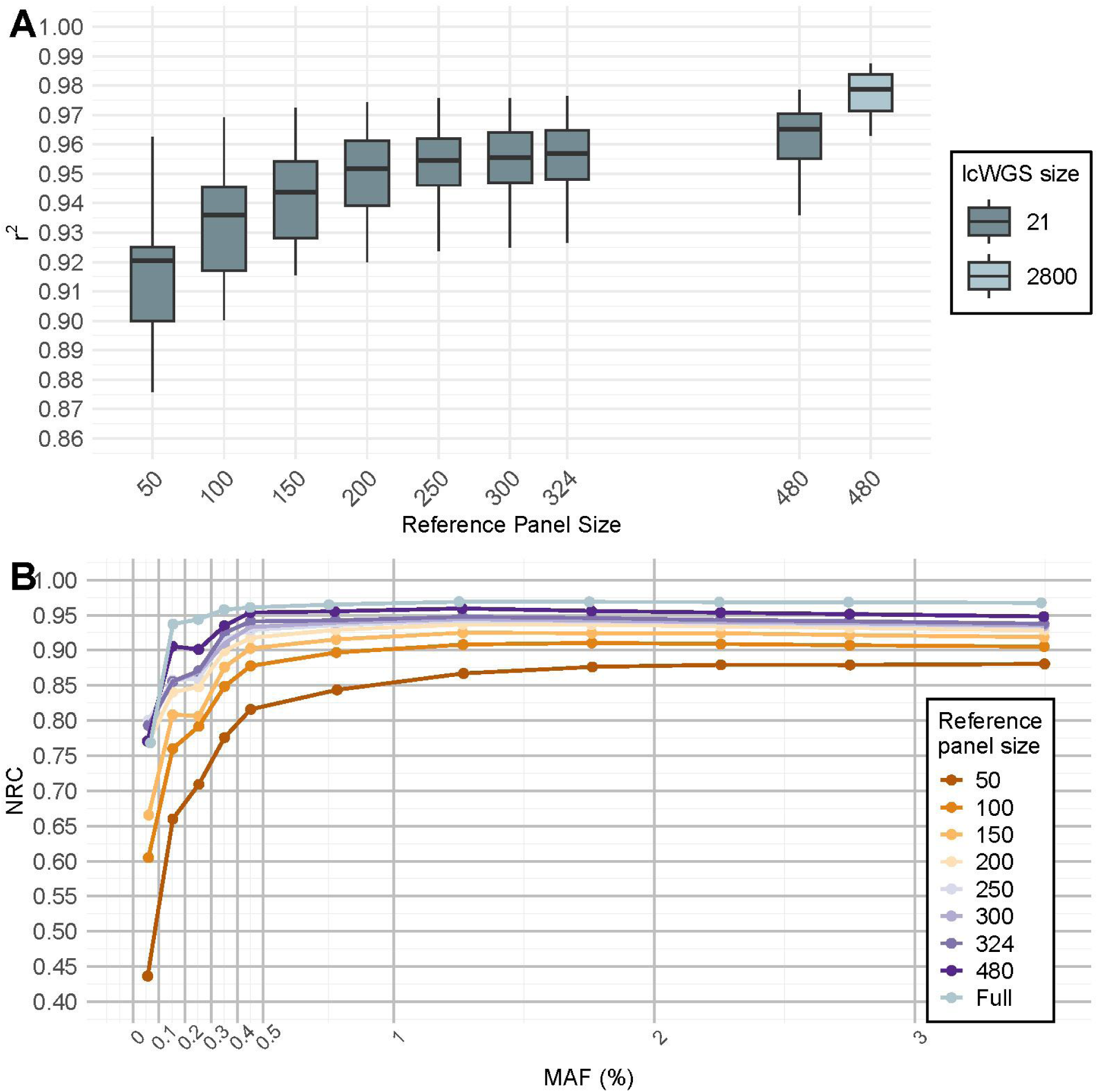
The effect of reference panel size on imputation accuracy. A) The per sample imputation accuracy using different reference panel sizes. 480 is the full reference panel size excluding the replicate individuals. B) The non-reference concordance (NRC) across minor allele frequency (MAF) bins for different reference panel sizes. ‘Full’ refers to the full reference panel of 480 samples along with 2800 lcWGS.

### The effect of lcWGS sample size

We tested imputation accuracy in subsets of the lcWGS without using a reference panel. We subsampled the lcWGS dataset, always including the 21 replicates, down to a size of 50,100,200,500,1000 and 2000 individuals. We chose these individuals either by sampling randomly across our dataset or by sampling full families, representing two realistic sampling schemes that might be encountered in a true study system and identified variants de-novo (number of variants per subset can be found in Supplementary Figure 16). We then imputed these datasets using STITCH. The accuracy of imputation benefited from increasing sample size (Figure 4A) with diminishing returns in accuracy gain when using more than 500 samples, which achieved an imputation accuracy of approximately 0.95 on average.

**Figure 4.**
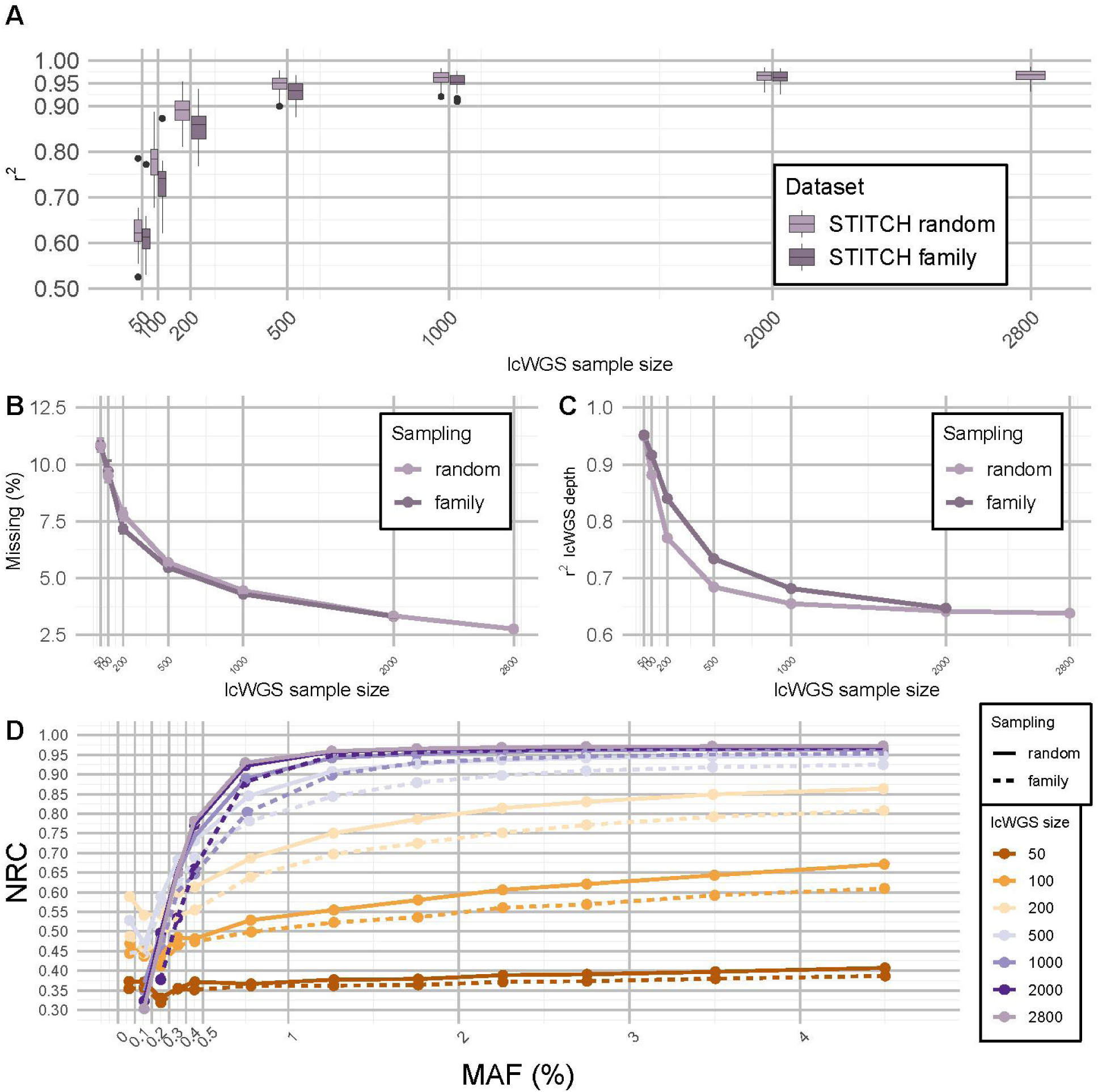
The effect of low-coverage sample size on imputation accuracy. A) The imputation accuracy per sample in the 21 replicates using a different number of low-coverage whole genome sequences (lcWGS) in STITCH. B) The average genotype missingness (95% CI shown as error bars) in genotypes imputed by STITCH across all samples in each subset. C) The correlation of imputation accuracy and lcWGS depth for the 21 replicates for different sizes of the lcWGS dataset, a larger correlation shows larger dependence on the underlying sequencing depth of each sample. D) The non-reference concordance across minor allele frequency bins for different lcWGS sizes. For all panels the subsampled dataset was generated by either random sampling or by sampling whole families of individuals.

Since we observed missing data and a dependence of per-sample imputation accuracy on lcWGS sequencing depth we quantified these aspects in smaller lcWGS sample sizes. The percentage of missing data decreased with increasing sample size, even after reaching 1000 samples (Figure 4B). To measure the dependence on lcWGS depth we estimated the correlation of imputation accuracy and lcWGS. A higher correlation implies a larger effect of lcWGS depth on performance. The correlation for small sample sizes (n=50, 100) was almost complete (>0.9) implying that imputation was rather ineffective at these sample sizes, and most genotypes resulted from observed data (Figure 4C). On the other hand above 500-1000 samples this metric, as overall imputation achieved its smallest observed value. Lastly, the non-reference concordance across allele frequency bins increased with increasing sample size and was very similar from 1000 samples and practically indistinguishable between 2000 and 2800 samples at 2x depth (Figure 4D). In all comparisons, a random subset of individuals performed better than a family stratified one which can be explained from the stochastic inclusion of more relatives of the replicate individuals in the random dataset (Supplementary Figure 17).

### Impact on downstream analyses

The homozygosity along the genome of an individual is a useful metric that can provide important insights such as the inbreeding state of a wild population with implications for conservation. In addition, because imputation with or without a reference panel performed differently when it came to inferring heterozygous genotypes, there may be an impact on the inference of runs of homozygosity (ROH) per individual. To address this question, we annotated homozygous-by-descent (HBD, equivalent to ROH) regions along the genome using the R package RzooRoH (Bertrand et al. 2019). The overall distribution of the ROH segments was very similar between datasets, with GLIMPSE identifying slightly more short ROH segments where STITCH does not (example region in Figure 5A). One major difference was the assignment of ROH segments into different length classes corresponding to a different age of the common ancestor. Specifically, using the genotypes imputed with STITCH, RzooRoH assigned more segments into younger classes (age < 32 generations ago) compared to the GLIMPSE dataset (Figure 5B). As a consequence, when calculating genome-wide inbreeding coefficients using only young ROH segments (< 32 generations ago) the results differed between the two methods (Figure 5C). In contrast, including older ROH classes resulted in equivalent inbreeding coefficients between the two methods (Figure 5C).

**Figure 5.**
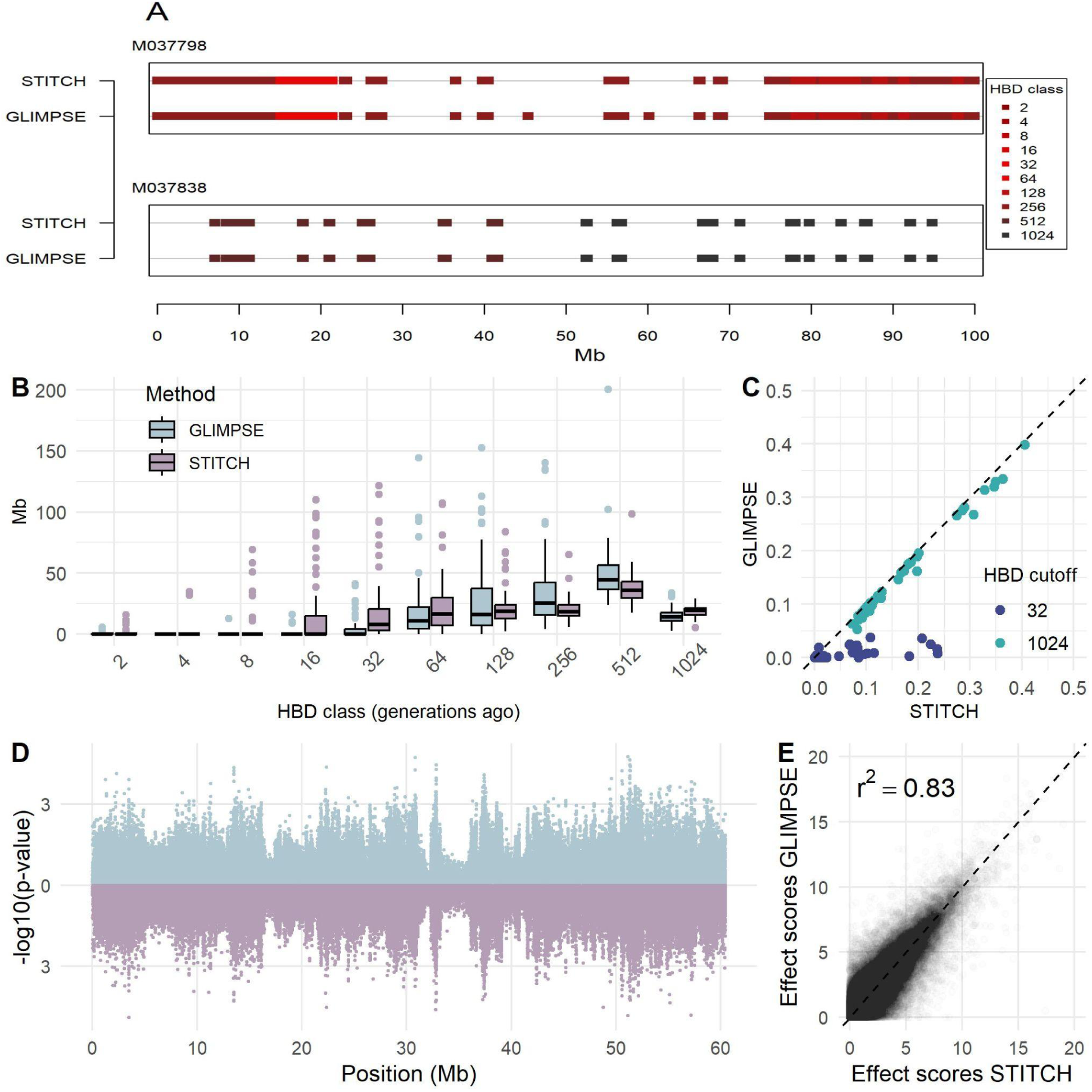
Downstream analyses in the imputed datasets. A) The homozygous-by-descent (HBD) segments inferred for an inbred (M037798) and a non-inbred sample (M037838) along the first 100Mb of the genome. HBD classes correspond to the two homozygous segments sharing a common ancestor that many generations ago.. B) The distribution of the length of all HBD segments assigned to a specific class using the 51 samples when imputed with reference panel (GLIMPSE) and when imputed only with lcWGS (STITCH). C) The proportion of the genome annotated in being HBD up to a certain number of generations ago between the two imputed datasets for the 51 individuals. D) Manhattan plot of -log10 p values from a GWAS on adult tarsus length with either imputed dataset (GLIMPSE above, STITCH below) for an example linkage group (LG2) E) Plot of effect scores among the two methods for the same SNPs as the p-values in panel D.

**Figure 6.**
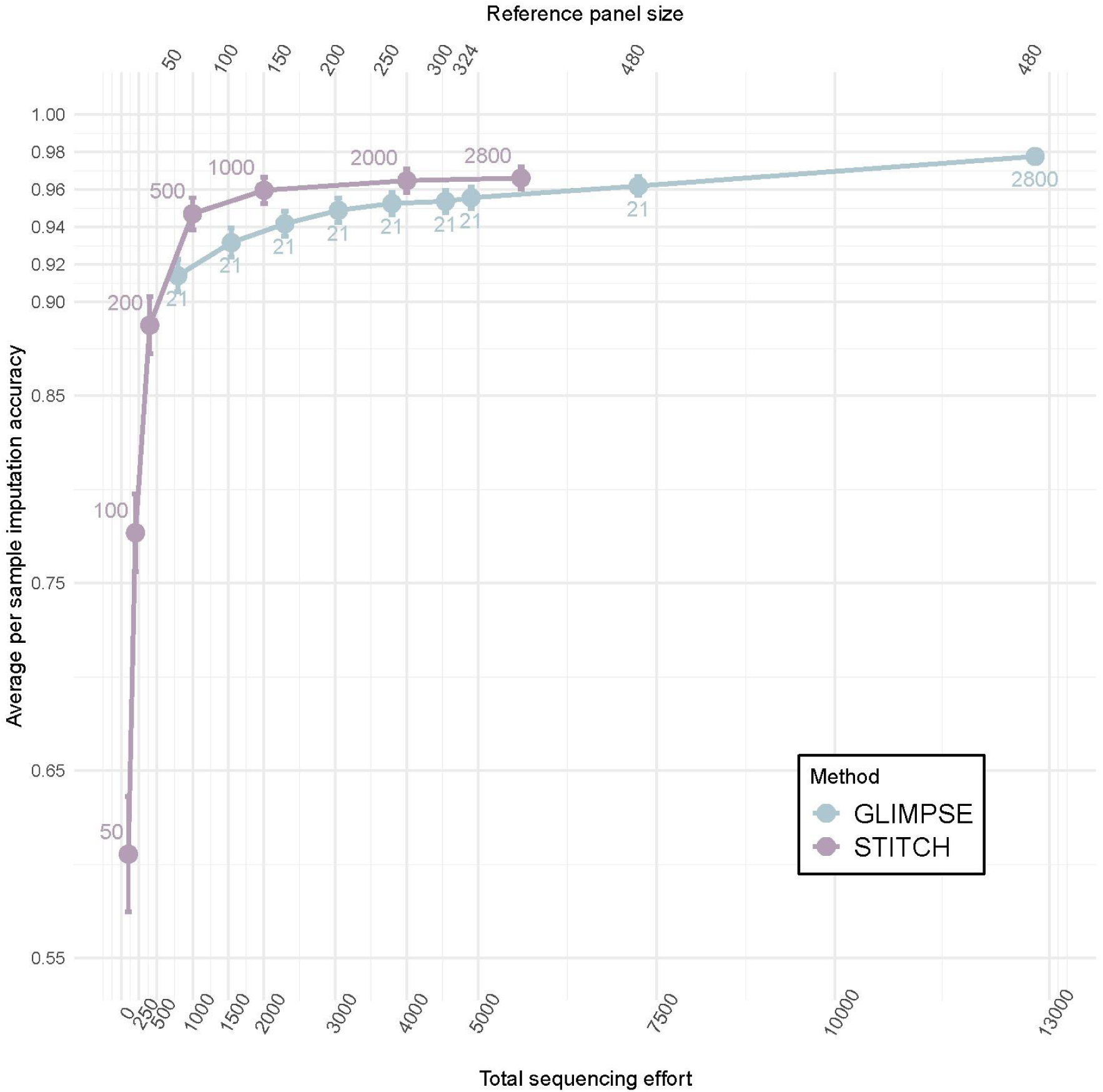
Comparison of imputation accuracy as a function of total sequencing effort. The imputation accuracy for all datasets in this study. Bottom x-axis is total sequence effort defined as 2 times the lcWGS size + 15 times reference panel size, with multipliers corresponding to average sequencing depth in each dataset. Top x-axis shows the reference panel size, shown only for GLIMPSE. Text by points shows the 2x lcWGS sample size imputed. Mean and 95% confidence intervals are estimated from the 21 replicate individuals.

One of the most frequent uses of whole genome sequences is to identify the regions of the genome affecting a phenotype (the trait’s architecture) using a genome-wide association study (GWAS). To test the performance of the different imputation methods on a GWAS, we looked for an association of the imputed genotypes with the length of the left tarsus, a trait with an intermediate heritability in adult birds (0.216 Hewett et al. 2025)). We selected only individuals with the phenotype measured at the adult stage of their life (N = 1980) and fitted their sex as a fixed effect. For the two imputed datasets we estimated the allele sharing genomic relationship matrix and fitted it as a random effect in a linear mixed model to account for confounding population and family structure. The results were very similar between GLIMPSE and STITCH and both the inferred effect scores and p-values were highly correlated (Squared pearson’s correlation for effect scores = 0.83 ; p-values = 0.73; an example linkage group can be seen Figure 5D, 5E). Full results in Supplementary Figure 18.

## Discussion

Low-coverage whole-genome sequencing (lcWGS) followed by imputation offers a cost-effective strategy for generating high-quality genotypes in large datasets. Traditional imputation methods rely on (an expensive) reference panel, but alternative approaches can perform imputation without a reference panel. In this study, we compare these two methods to establish best practices for imputation in wild populations. Using 21 samples sequenced in both low and high coverage as a validation set, we demonstrate that imputation of 2,800 lcWGS samples is highly accurate when a reference panel of 502 individuals is used. Reducing the reference panel size to 50 individuals from the focal population still achieves accuracy larger than 90% and increasing either the reference panel size or the lcWGS dataset improves performance. In the absence of a reference panel, imputation of the 2,800 lcWGS with STITCH produces comparable accuracy to GLIMPSE, though it struggles with low-frequency alleles (<1%). For smaller lcWGS data sets, imputation accuracy can be high from 500 samples, provided rigorous quality control is applied. When using the entire lcWGS dataset, a polygenic GWAS revealed minimal differences between imputations with or without a reference panel, but some caution was required when looking into runs of homozygosity (ROH). We conclude with recommendations for using imputation in lcWGS studies of wild populations.

### Imputation with a reference panel

Imputation using a reference panel has been shown to be very accurate in humans, domestic animals, crop plants, and simulated data (Swarts et al. 2014; Das et al. 2018; Deng et al. 2022; Teng et al. 2022). Recently it has been demonstrated that it performs well in wild populations (Watowich et al. 2023; Tan et al. 2025). Watowich et al., (2023) quantified imputation accuracy in two empirical datasets of different species, one with a large reference panel (n=741; rhesus macaques *Macaca mulatta*) and one with a small reference panel (n=68; gelada monkeys *Theropithecus gelada*). The authors found accurate imputation in both datasets, even for the small reference panel sizes in the gelada dataset (r^2^ = 0.87 and 0.96 for allele frequencies in the range of 1-5% and 10-50% respectively). In addition Tan et al. 2025, attained very high imputation accuracy by using a limited number of individuals as a reference panel (n=29) (>0.9) in the hihi/stitchbird (*Notiomystis cincta*). However, the gelada dataset harbours a small number of polymorphic sites, low heterozygosity and elevated ROH (Chiou et al. 2022; Watowich et al. 2023) and the hihi has notoriously low diversity due to a dramatic past bottleneck event leaving approximately half of the genome homozygous by descent (de Villemereuil et al. 2019; Duntsch et al. 2023). The barn owl dataset used here displays a larger diversity (heterozygosity ∼ 0.2, approx. 1 SNP / 100bp) due to a well-connected continental population and rare occurrence of inbreeding (Cumer, Machado, Dumont, et al. 2022; Hewett et al. 2025). It also exhibits linkage disequilibrium (LD) patterns characteristic of birds, with microchromosomes of elevated recombination rates (low LD) and large chromosomes of diverse recombination landscapes (Topaloudis et al. 2025). Therefore, it represents a distinct parameter space of the diversity and LD values studied so far. Based on our investigation we show that imputation using a large reference panel (502 individuals) coupled with 2X lcWGS can generate imputation with accuracy greater than 0.97 (r^2^ > 0.97) even for a species with genetic diversity larger than those studied previously and this result can extend to low-frequency alleles (MAF > 0.2%).

An important consideration for non-model species where resources are limited is the size of the reference panel for accurate imputation results. We found high imputation accuracy (r^2^ > 0.9) even when reducing the size of the reference panel to 50 individuals, and while any increase beyond 50 samples increased the resulting accuracy, diminishing returns were observed above 200 individuals. While imputation of rare alleles performed worse with decreasing reference panel size, this result is likely irrelevant since in low sample sizes very rare alleles will be absent due to sampling variance. For example, filtering out alleles present in less than two copies, a lenient filtering threshold, applies a 2,1,0.5% MAF filter in a reference panel of 50, 100 and 200 individuals respectively. Thus, we showed that despite substantial genetic diversity in the barn owl, we also find accurate imputation with a reference panel of 50 individuals (Watowich et al. 2023; Tan et al. 2025), a promising result for studies in non-model species.

Concerning the composition of the reference panel, despite a large number of individuals (n=346) of the focal Swiss population, there was improved imputation accuracy when including distantly related samples. This result is in line with studies in humans (Purnomo et al. 2025), dairy cattle (Lloret-Villas et al. 2023) and Nile tilapia (*Oreochromis niloticus*) (Garcia et al. 2022), where the inclusion of a large and diverse reference panel outperforms population-specific panels which lack an exhaustive representation of the haplotypes present in the target population (but see (Dekeyser et al. 2023) for an investigation into the possible caveats of this practice). Similar results are found by studies in aDNA, where relevant reference panels might be impossible to collect (Sousa da Mota et al. 2023; Bougiouri et al. 2024; Erven et al. 2024). Our results therefore support the conclusion that even in the presence of a genetically relevant reference panel, combining all available resources can improve the imputation accuracy.

A less discussed factor influencing imputation accuracy is the size of the lcWGS dataset. Because GLIMPSE uses the imputed lcWGS haplotypes as part of the reference panel (Rubinacci et al. 2021), increasing the lcWGS dataset can also provide an increase in imputation accuracy. We observed a 1.6% increase on average when imputing the full set of 2800 lcWGS versus imputing only the 21 lcWGS replicates using the full reference panel. This behaviour of GLIMPSE can lead to an underestimation of imputation accuracy when the target lcWGS sample size is small, for example in leave-one-out settings of estimating imputation accuracy and should be accounted for.

Lastly, an important observation was the impact of sequencing depth of the ‘truth set’ in the resulting imputation accuracy. Through our diverse set of replicate individuals we found that samples sequenced at intermediate depth (< 20x) in the truth set, showed a lower imputation accuracy than samples sequenced at higher depth (> 20x). This is due to a reduction of the heterozygous genotypes in the ‘truth set’, a well known caveat of low to medium sequencing depth (Nielsen et al. 2011; Kardos and Waples 2024). Interestingly, these heterozygotes were inferred in the imputed dataset showing that imputation with a reference panel can outperform genotype calling at intermediate sequencing depth. Based on this finding, we suggest ideally using the individuals sequenced at highest depth (>20x) in each study to validate the imputation results, and build a reference panel. Failure to do so might underestimate the accuracy of imputation.

Assessing imputation accuracy using a real dataset can incorporate realistic variation and errors in the data generating process (library preparation, sequencing depth, mapping etc.) but explores a parameter space defined by the particularities of the study system. What characterises our dataset is a high degree of relatedness among individuals of the reference panel, the lcWGS dataset and between the two. High relatedness in the reference panel can improve haplotype phasing (Glusman et al. 2014) but will decrease the number of observed haplotypes, reducing the ‘effective sample size’ which might explain why including non-Swiss samples marginally improved imputation accuracy. Relatedness within the lcWGS dataset can provide an increase in imputation accuracy when using a large lcWGS dataset. Indeed such a structure might be responsible for the high imputation accuracy seen when using the full lcWGS dataset instead of only the 21 replicates. Lastly, relatedness between the lcWGS and the reference panel can similarly boost imputation accuracy by long haplotype sharing among close relatives. The latter might be partially responsible for the very accurate imputation of rare alleles, a behaviour expected when using pedigree-based imputation methods (Liu et al. 2019) but here observed using GLIMPSE, a linkage-disequilibrium (LD) based method. In summary, while some aspects of our results might be influenced by the specificities of our study species, the broad findings should hold for all wild species. In addition, in long-term monitoring studies where similar familial structures might emerge, we illustrate the benefits of imputation even for low-frequency alleles, when using a reference panel with standard LD-based imputation methods.

### Imputation without a reference panel

Imputation of lcWGS using STITCH can be a competitive alternative when a reference panel is not available, although imputation accuracy will depend on the size and the depth of the lcWGS dataset. In past studies as well as in our own results, there appears to be a “critical size” below which genotype refinement is mostly inaccurate, and above which diminishing returns are observed. Importantly, this critical size should be dependent on the underlying genetic diversity of the organism and the lcWGS sequencing depth. In species with low genetic diversity, maximum accuracy can be achieved with smaller sizes and sequencing depths, (e.g. 200 dairy cattle at 1x (Teng et al. 2022), 500 CFW mice at 0.1x (Davies et al. 2016), 120 Pacific oysters (*Crassostrea gigas*) at 1x (Yang et al. 2024)). On the contrary, in more genetically diverse species the sample size and sequencing depth required to achieve similar imputation quality is often larger (e.g. 500 owls at 2x (this study), at least 1000 human samples at 1x (Davies et al. 2016), and 500 samples at 2x in a simulation study with intermediate diversity and LD (Lou et al. 2021)). Past comparisons show that the maximum attainable accuracy can be limited by lcWGS depth, meaning that a lcWGS dataset of 0.5x might never achieve the same imputation accuracy of a dataset in 2x regardless of sample size (Davies et al. 2016; Lou et al. 2021; Yang et al. 2024). In our investigation, variation in sequencing depth (a realistic expectation in any sequencing experiment) resulted in varying coverage for our replicate samples. We found individuals with a lower coverage depth had a smaller imputation accuracy, an effect which becomes more pronounced with fewer individuals sequenced. Therefore, our results prove that imputation without a reference panel can be accurate, provided that the dataset meets minimum requirements in number of samples and sequencing depth, which in our case were > 500 and 2x, respectively.

Despite its overall accuracy, imputation using only lcWGS had two main shortcomings: poor performance for rare alleles and a proportion of missing data. Lower performance in rare alleles (MAF < 1%) is an expected outcome of imputation without a large reference panel and while filtering out rare alleles might introduce biases (e.g. in kinship estimation (Weir and Goudet 2017)) it is common practice, especially for GWAS where power is low when alleles are rare. On the contrary, the missing data is an intended behaviour using default parameters in STITCH which does not call genotypes with a genotype probability smaller than 0.9. This behaviour has spurred previous studies to suggest imputing the sporadic missing data after STITCH, i.e. using BEAGLE (Teng et al. 2022; Vi et al. 2025). Here we show that missing genotypes can be inferred from genotype probabilities with minimal accuracy loss eliminating the necessity for a second imputation step. Notably, both missingness and poor imputation in low frequency alleles are exaggerated in small lcWGS datasets so studies planning to use a small lcWGS dataset without a reference panel should address these issues. We assume a valid solution to be a relevant MAF filter (e.g. 1-2%) and using a more lenient genotype probability threshold (e.g. 0.8), or the genotype likelihoods themselves, if missing data is an issue.

### Downstream analyses

In downstream analyses STITCH performed similar to GLIMPSE in a polygenic trait GWAS. In the GWAS of tarsus length, estimated effects were highly correlated between the two methods. In general, a dataset imputed with an imputation accuracy *r*^2^ on *N* samples will have equal power as a dataset with *r*^2^ × *N* samples where all genotypes are known (Das et al. 2018).

Thus, given the similar imputation accuracy estimates among the two approaches, the power of the two datasets is similar and high agreement in the GWAS is to be expected. Therefore, imputation of lcWGS without a reference panel can be a powerful approach to identify the genetic architecture of traits in wild species (Hooper et al. 2024; Justen et al. 2024).

For calling runs of homozygosity (ROH), there were subtle differences between the two imputation methods. Past studies show that imputation with a reference panel can infer ROH in agreement with high coverage data (Sousa da Mota et al. 2023; Bougiouri et al. 2024; Erven et al. 2024). We further show that regardless of whether a reference panel was present or absent, regions inferred as ROH were similar, although imputation without a reference panel inferred a larger proportion of young ROH segments. This behaviour could be due to differences between the two datasets, such as the estimated allele frequencies in the 51 samples due to misimputed alleles or missing data. Another difference might be a more extensive set of variants considered in the absence of a reference panel, where variant discovery was performed in 2800 samples, contrary to the reference panel of 502 samples. Thus, when inferring ROH in an imputed dataset, some variation in the fine scale (length of ROH segment, age of segment) might be expected due to methodological differences between imputation methods.

### Implications & Perspectives

An understanding of the strengths and limitations of different imputation strategies can help balance a fixed budget with the objectives of a study. Using our results we can test the imputation quality of different combinations of sample sizes in the absence or presence of a reference panel (Supplementary Figures 19-20). Imputation without a reference panel with at least 500 lcWGS samples in 2x, returned an imputation accuracy that was unmatched unless a reference panel of at least 200 samples was used. Therefore, for an experiment generating genotypes for at least 500 lcWGS sequences at 2x, the addition of a reference panel of up to 200 samples will only bring marginal benefits. On the other hand, imputation with lcWGS is unreliable in sample sizes below 500 samples. Ultimately, the best quality imputation, especially in rare alleles, can only come in the presence of a large relevant reference panel. We note that our results may underestimate the accuracy of GLIMPSE, especially because of the small lcWGS sample size tested, but can serve to illustrate that lcWGS without a reference panel has a place in the generation of large datasets in non-model species.

Many aspects of imputation in wild datasets remain to be explored as genomic imputation gains traction in non-model systems. The reality might be that every investigation will be served by different combinations of underlying genetic diversity, sample relatedness, attainable reference panel size and affordable sequencing depth. Presently, there is a dearth of studies providing benchmarks of imputation accuracy in the different regions of the parameter space and assessing the impact in relevant downstream analyses. While many case studies in different species can reveal the limitations and benefits of different imputation approaches under different realistic conditions, an additional valuable resource might be the development of versatile and easy to implement benchmarking tools. For example, publicly available catalogs of species’ demographic and genetic parameters (Adrion et al. 2020; Gower et al. 2025) coupled with powerful simulation software (Kelleher et al. 2016) allow the generation of large datasets. Such resources coupled with much needed, accessible imputation pipelines (Das et al. 2016) to quickly and reproducibly implement genomic imputation will allow researchers to test imputation performance under a feasible parameter space of reference panel sizes, lcWGS sample sizes, demographies and genetic diversity (Vi et al. 2025). Benchmarking in real and simulated data has benefits and drawbacks, and both will be useful in providing accurate guidelines for sound implementation of imputation in wild systems.

## Conclusion

We have tested the performance of imputing low-coverage (2x) whole genome sequences with and without a reference panel. Imputation with a reference panel is increasingly being used in studies of wild species and we have demonstrated very accurate imputations in a pedigreed population even for allele frequencies below 1%. High imputation accuracy (r^2^ > 0.9) can be achieved with reference panel sizes as low as 50 individuals. In the absence of a reference panel, lcWGS at 2x with genotype refinement, coupled with reasonable quality control, can be a viable alternative with 500 samples or more. We hope that accurate imputation methods, careful benchmarking, and increasingly affordable DNA sequencing will usher wild populations into the era of large genomic datasets.

## Acknowledgements

The authors want to thank Simone Rubinacci for discussions before data generation and helpful feedback during imputation using GLIMPSE. They also thank the Lausanne Genomic Technologies Facility (GTF) and specifically Julien Marquis and Melanie Dupasquier for aid during library preparation and data generation.

## Author contributions

BA and AR maintain the long term barn owl study system. ALD and CS performed DNA extraction, and library preparation. AT, TC and JG conceived and planned the study with help from OD. AT, TC and EL imputed the sequencing data. AT performed comparisons and downstream analyses with help from TC and AH and EL. AT wrote the manuscript with help from all authors.

## Data availability

Sequence data used in the study from previous publications are available on NCBI under BioProject codes PRJNA700797, PRJNA727915, PRJNA727977, PRJNA774943, PRJNA925445 and PRJNA1172395. Low coverage sequencing data can be found under BioProject code XXXXXX. Scripts available at https://github.com/topalw/Imputation_owls. Downstream data will be made available pre-publication.

