## Supplementary Material for "Benchmarking imputation accuracy in the presence or absence of a reference panel"

### Supplementary Text 1

#### Imputation using BEAGLE

We genotyped individuals sequenced at low coverage using a combination of ANGSD v.0.931 (Korneliussen et al. 2014) for SNP calling followed by Beagle v.3.3.2 (Browning and Browning 2009) for phasing and imputation, following a pipeline similar to that implemented in (Lou et al. 2021).

Genotypes and genotype dosages were first estimated in ANGSD using the following settings: *-GL 2 -doGlf 2 -doMaf 1 -doMajorMinor 4 -doCounts 1 -remove_bads 1 -minMapQ 30 -minQ 20 -skipTriallelic 1 -minInd 1000 -setMaxDepthInd 10 -setMinDepthInd 1 -SNP_pval 1e-6.*

SNP calling was restricted to regions of the genome with high mappability, as described in Corval et al. (2023). Genotype imputation was then performed using Beagle v.3.3.2 with default parameters, using the genotype likelihoods output by ANGSD in Beagle format (.beagle.gz) as input. Imputation was conducted without a reference panel.

For the imputed datasets generated by BEAGLE we compared the dosage (rounded to integers) with the dosage of the high coverage dataset of the 21 replicates. We quantified performance the same way we did for STITCH. We also quantified the reduction in sample size by using the same set of individuals as in the STITCH subsets.

#### Supplementary Tables

**Supplementary Table 1. Number of SNPs used in the study. Percentages in parentheses are compared to the non filtered datasets.**

| Method | Number of SNPs | | |
|---|---|---|---|
| | **No filtering** | **0.6** | **0.8** |
| **GLIMPSE** | 10,102,233 (100%) | 9,709,936 (96%) | 9,526,404 (94%) |
| **STITCH** | 24,224,697 (100%) | 21,322,604 (88%) | 20,305,801 (84%) |
| **BEAGLE** | 45,746,186 (100%) | 17,303,660 (38%) | 8,212,527 (18%) |

**Supplementary Table 2. Number of unique and overlapping SNPs among methods. Method–unique SNPs in the diagonal. Proportion of each method's total (row) in parentheses.**

| Method | GLIMPSE | STITCH | BEAGLE |
|---|---|---|---|
| **GLIMPSE** | 533,585 (5%) | 8,065,481 (80%) | 9,567,325 (95%) |
| **STITCH** | 8,065,481 (80%) | 126,644 (0.5%) | 24,096,730 (99%) |
| **BEAGLE** | 9,567,325 (20%) | 24,096,730 (52%) | 20,160,403 (44%) |

#### Supplementary Figures

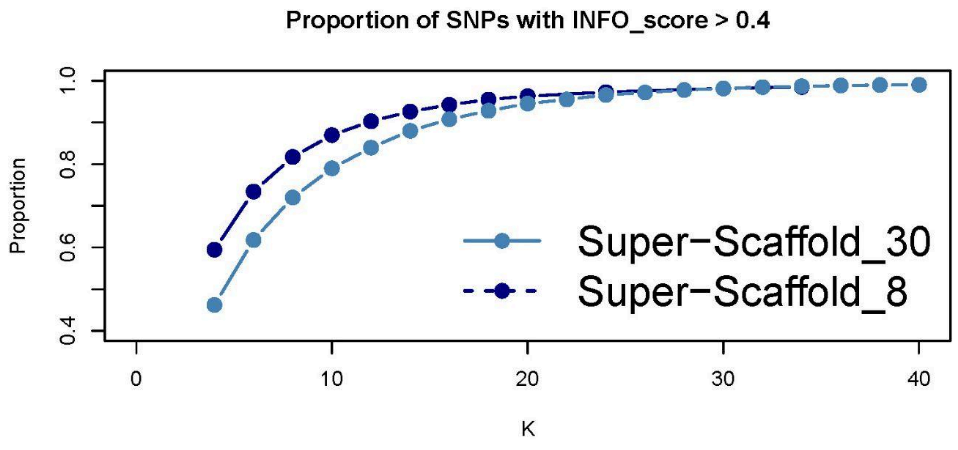

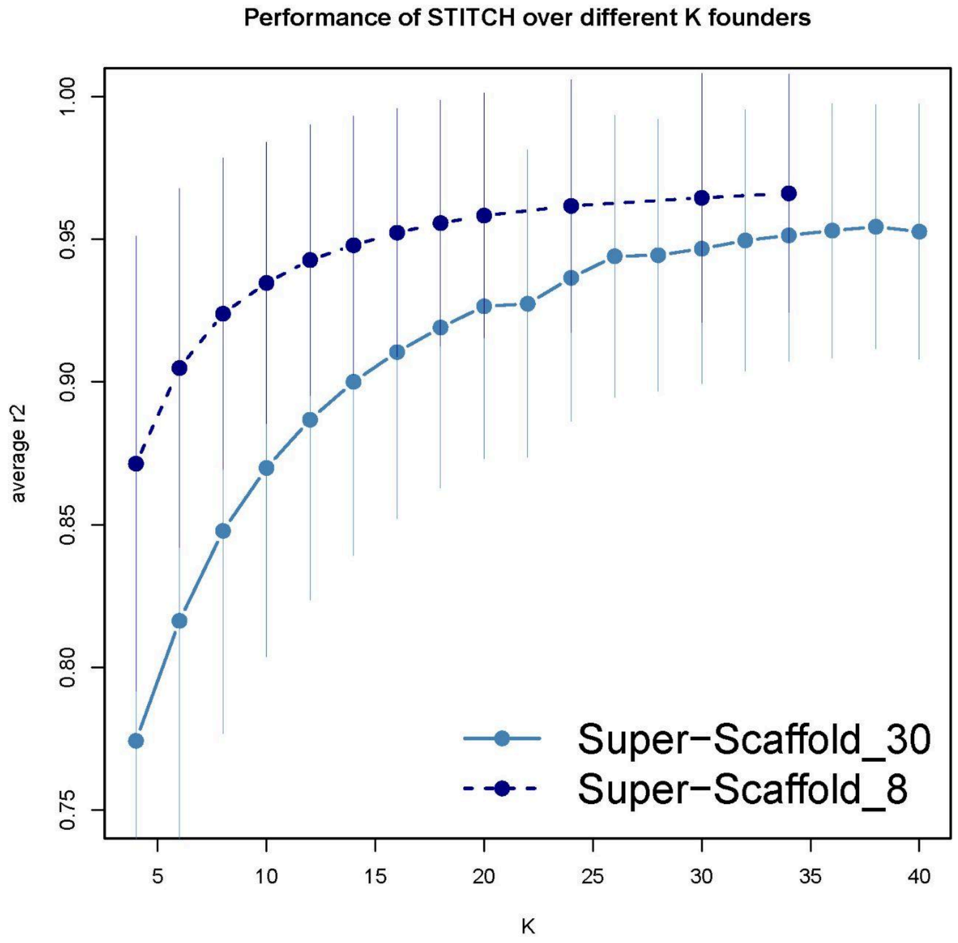

**Supplementary Figure 1. Choosing the right K in STITCH.**

Top panel shows the proportion of SNPs, relative to the total number of SNPs, with an information score larger than 0.4. The bottom panel shows the imputation accuracy of the 32 replicates for different values of K for each scaffold.

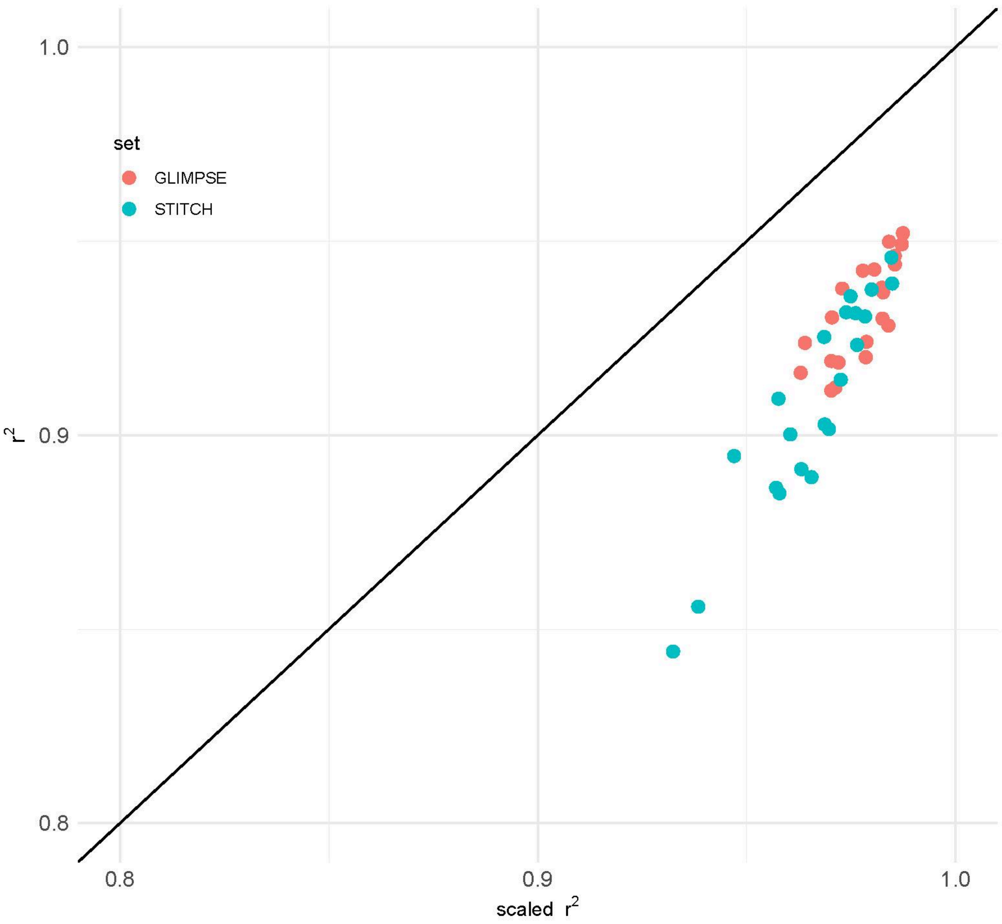

**Supplementary Figure 2 - Relationship between imputation accuracy and scaled imputation accuracy.**
Scaled imputation accuracy scales the dosage vector by the mean dosage and divides by the standard deviation. Scaled r2 gives more weight to rare alleles.

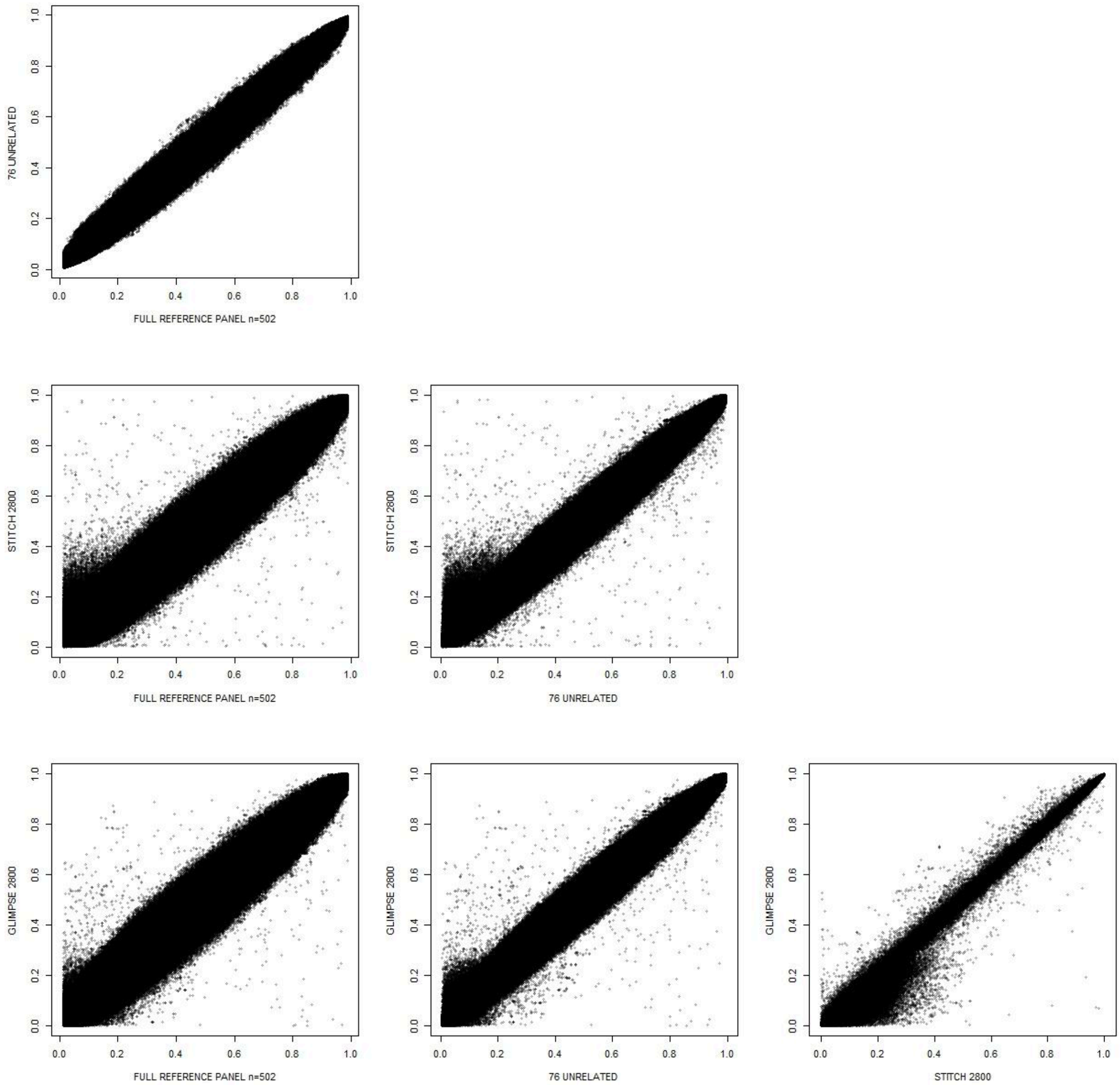

**Supplementary Figure 3. The comparison between allele frequencies estimated with different datasets**

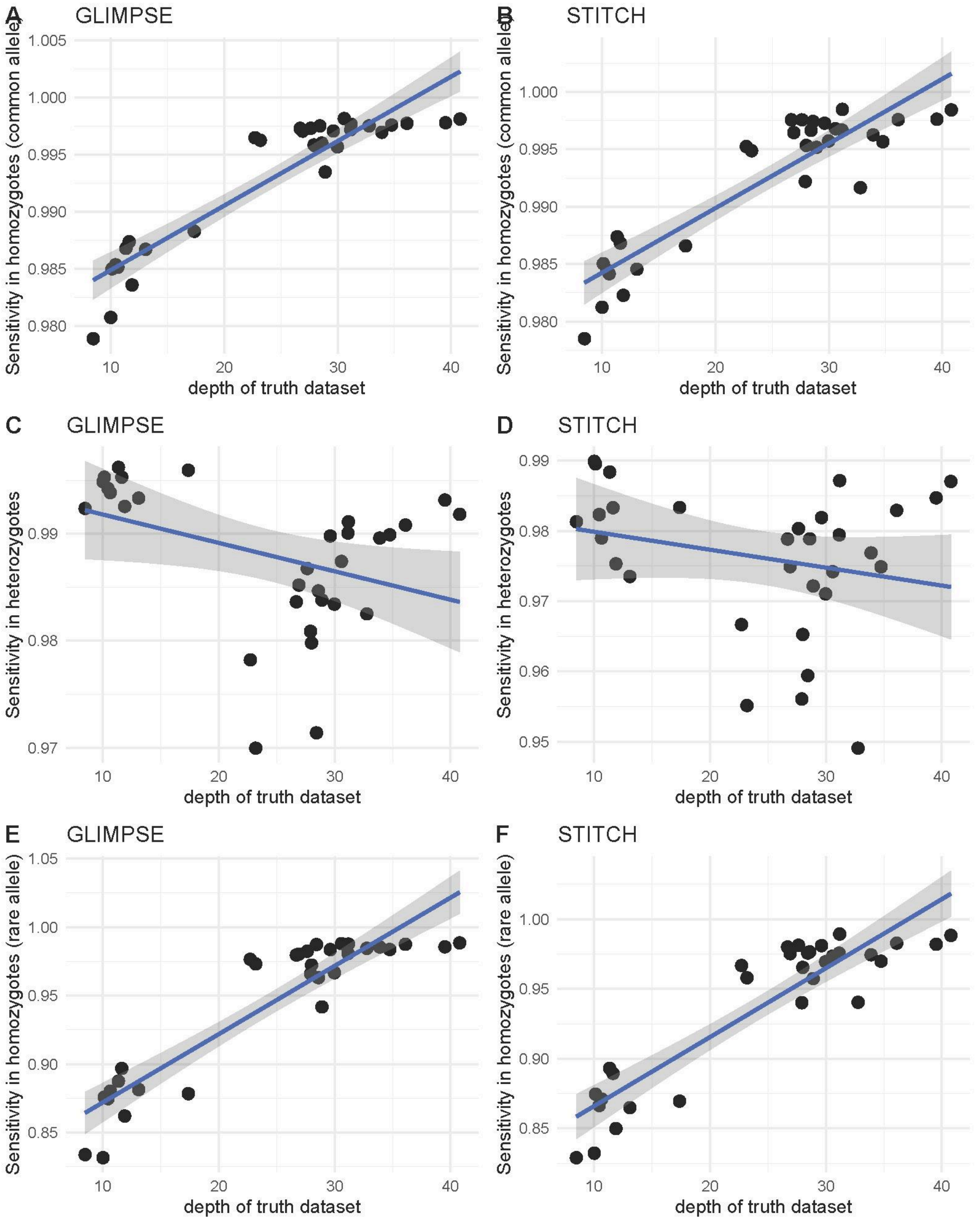

**Supplementary Figure 4. Sensitivity of each genotype class (rows), with each method (columns) as a function of the sequencing depth used as the 'truth'.**

Sensitivity refers to the ratio of true positives over true positives and false negatives and can be thought of as the proportion of true genotypes in one class (e.g. heterozygotes) inferred by the imputation method. **A,B**) Sensitivity of homozygotes for the common allele with GLIMPSE (A) and STITCH (B), **C,D**) Sensitivity of heterozygotes with GLIMPSE (C) and STITCH (D), **E,F**) Sensitivity of homozygotes for the rare allele with GLIMPSE (E) and STITCH (F). Note the different pattern among heterozygotes and homozygotes.

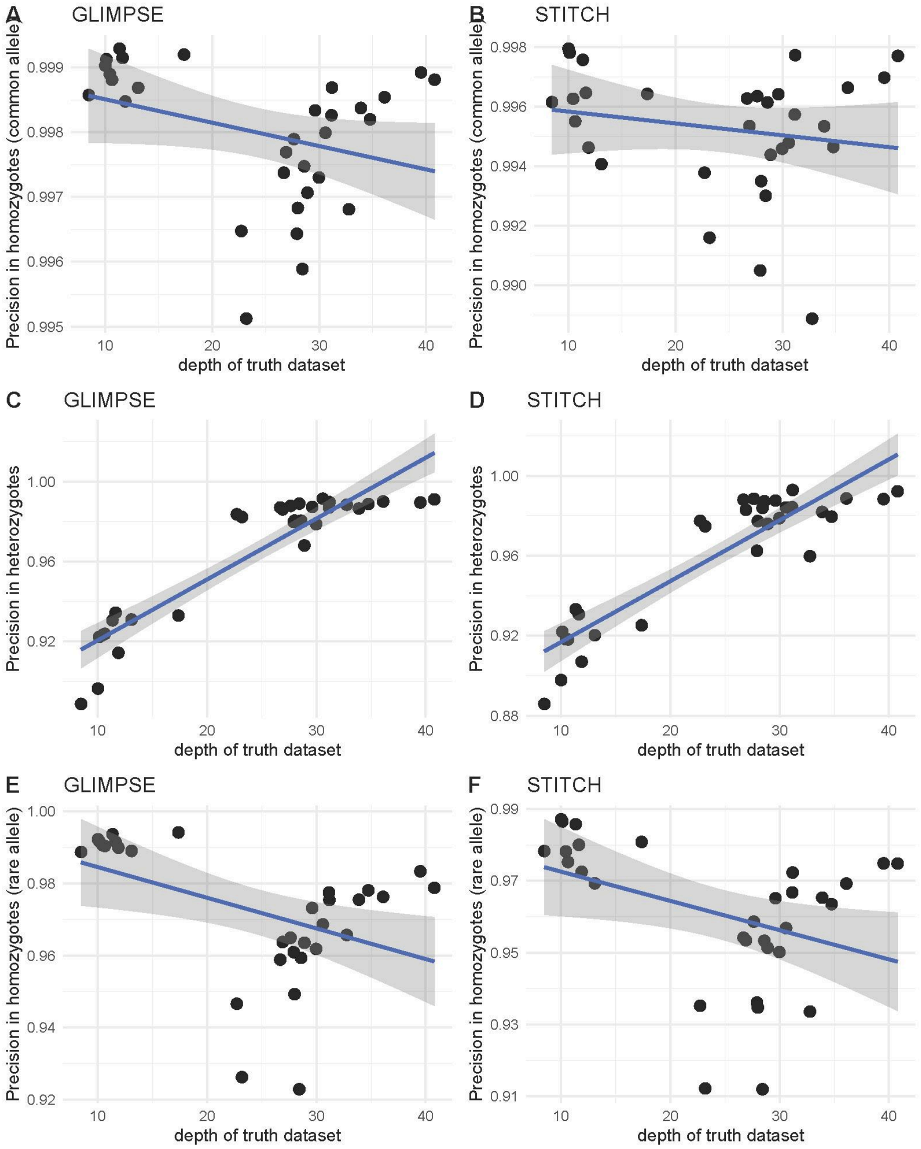

**Supplementary Figure 5. Precision of each genotype class (rows), with each method (columns) as a function of the sequencing depth used as the 'truth'.**

Precision refers to the ratio of true positives over true positives and false positives and can be thought of as the proportion of genotypes imputed in one class (e.g. heterozygotes) truly being in that class. **A,B**) Sensitivity of homozygotes for the common allele with GLIMPSE (A) and STITCH (B), **C,D**) Sensitivity of heterozygotes with GLIMPSE (C) and STITCH (D), **E,F**) Sensitivity of homozygotes for the rare allele with GLIMPSE (E) and STITCH (F). Note the different pattern among heterozygotes and either homozygotes.

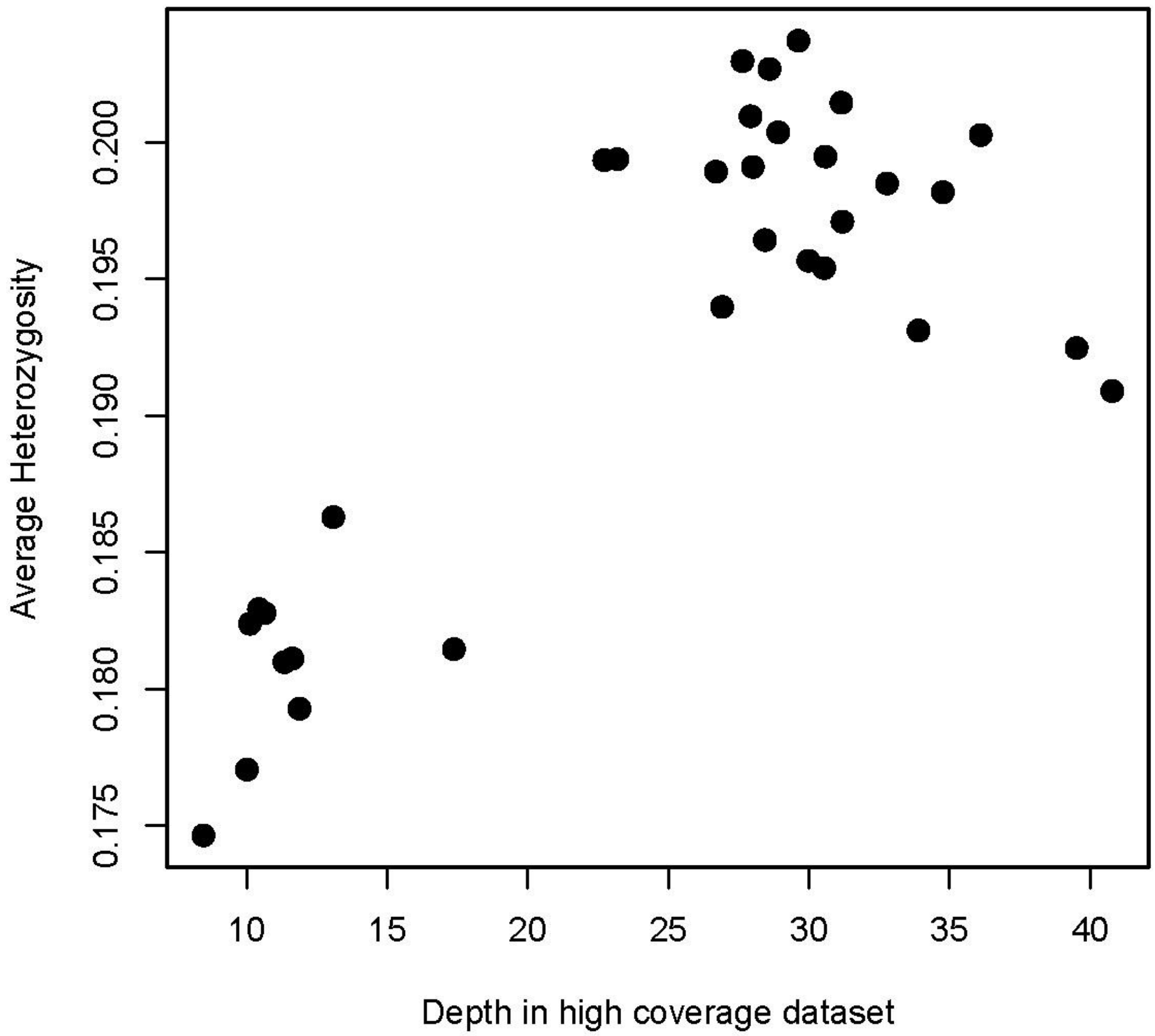

**Supplementary Figure 6. The effect of sequencing depth on heterozygosity.**

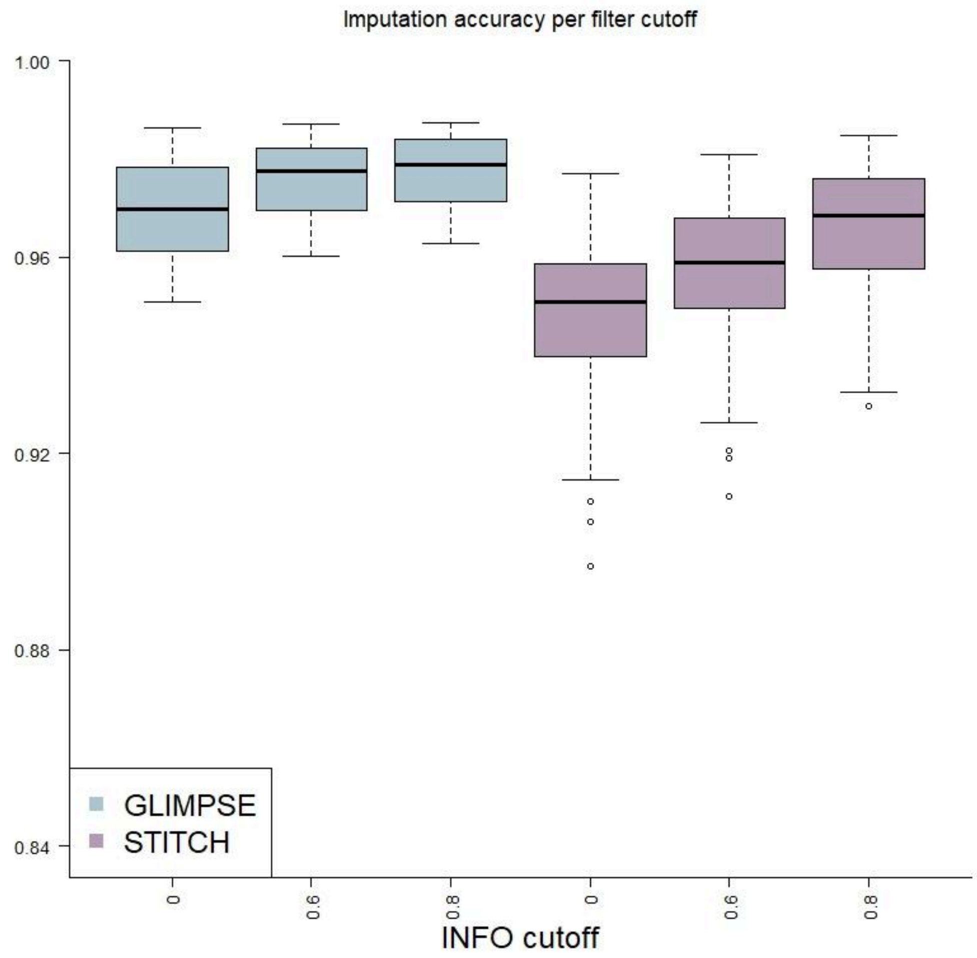

**Supplementary Figure 7 The effect of filtering cutoff in imputation accuracy.**

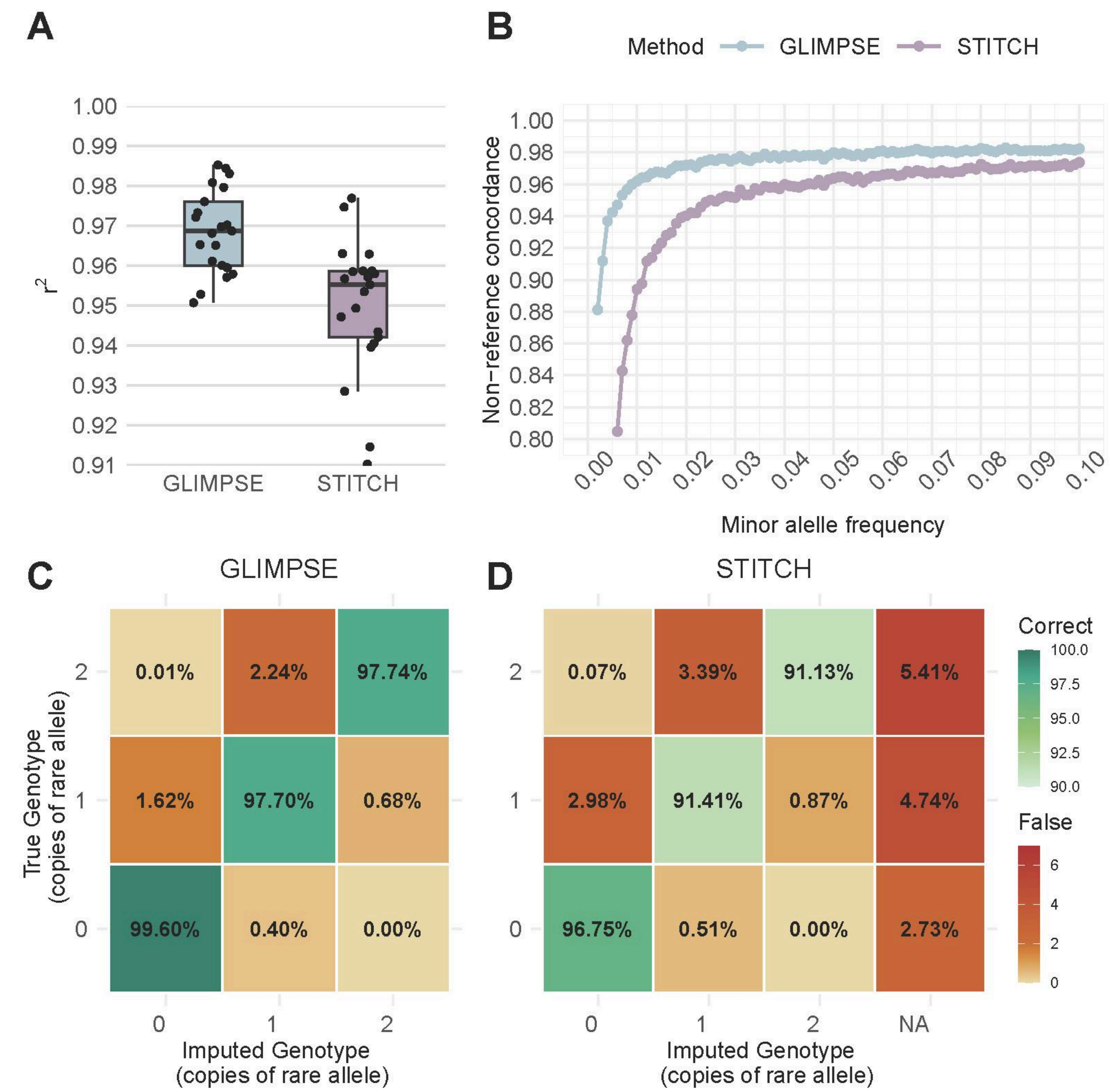

**Supplementary Figure 8 - The accuracy of imputation with no INFO cutoff.**

**A)** the squared correlation (imputation accuracy) per sample between the 'truth' (high-coverage data) and the dataset imputed by GLIMPSE or STITCH. Each point is one of the 21 replicate individuals**. B)** Non-reference concordance (NRC) per minor allele frequency bin. Each point represents the mean in the specific allele frequency bin. Allele frequency as defined in 76 unrelated individuals sequenced in high coverage. **C)** Confusion matrix for genotype classification for GLIMPSE. Each row represents the true genotype for these markers and the column the imputed one. **D)** Confusion matrix for genotype classification for STITCH. Each row represents the true genotype for these markers and the column the imputed one. In D the NA column represents uncalled genotypes in STITCH. For C and D, diagonals are colored differently since higher is better, than off-diagonal where lower is better. All statistics calculated along the whole autosomal genome

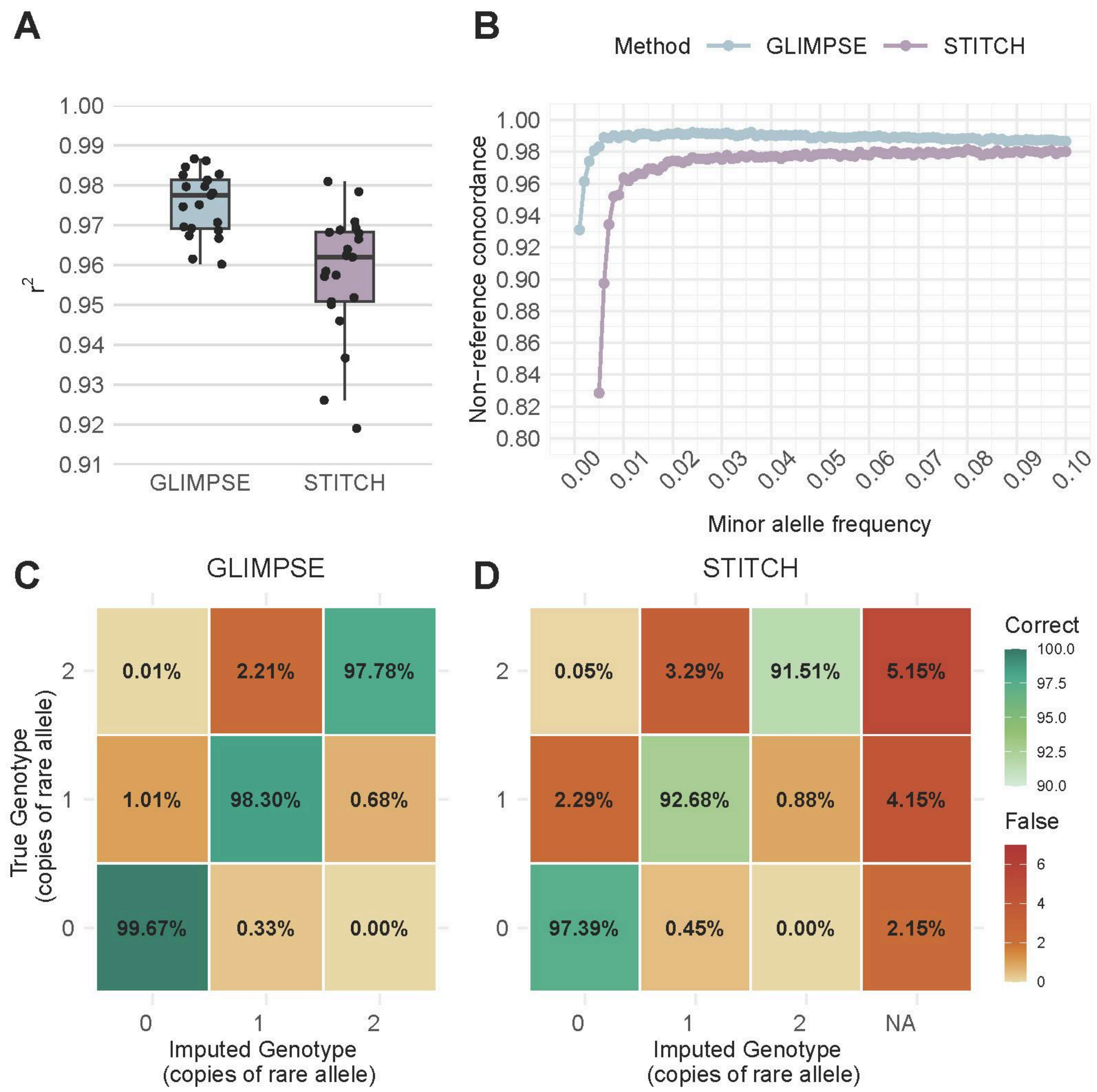

**Supplementary Figure 9 - The accuracy of imputation with an INFO cutoff of 0.6.**

**A)** the squared correlation (imputation accuracy) per sample between the 'truth' (high-coverage data) and the dataset imputed by GLIMPSE or STITCH. Each point is one of the 21 replicate individuals**. B)** Non-reference concordance (NRC) per minor allele frequency bin. Each point represents the mean in the specific allele frequency bin. Allele frequency as defined in 76 unrelated individuals sequenced in high coverage. **C)** Confusion matrix for genotype classification for GLIMPSE. Each row represents the true genotype for these markers and the column the imputed one. **D)** Confusion matrix for genotype classification for STITCH. Each row represents the true genotype for these markers and the column the imputed one. In D the NA column represents uncalled genotypes in STITCH. For C and D, diagonals are colored differently since higher is better, than off-diagonal where lower is better. All statistics calculated along the whole autosomal genome.

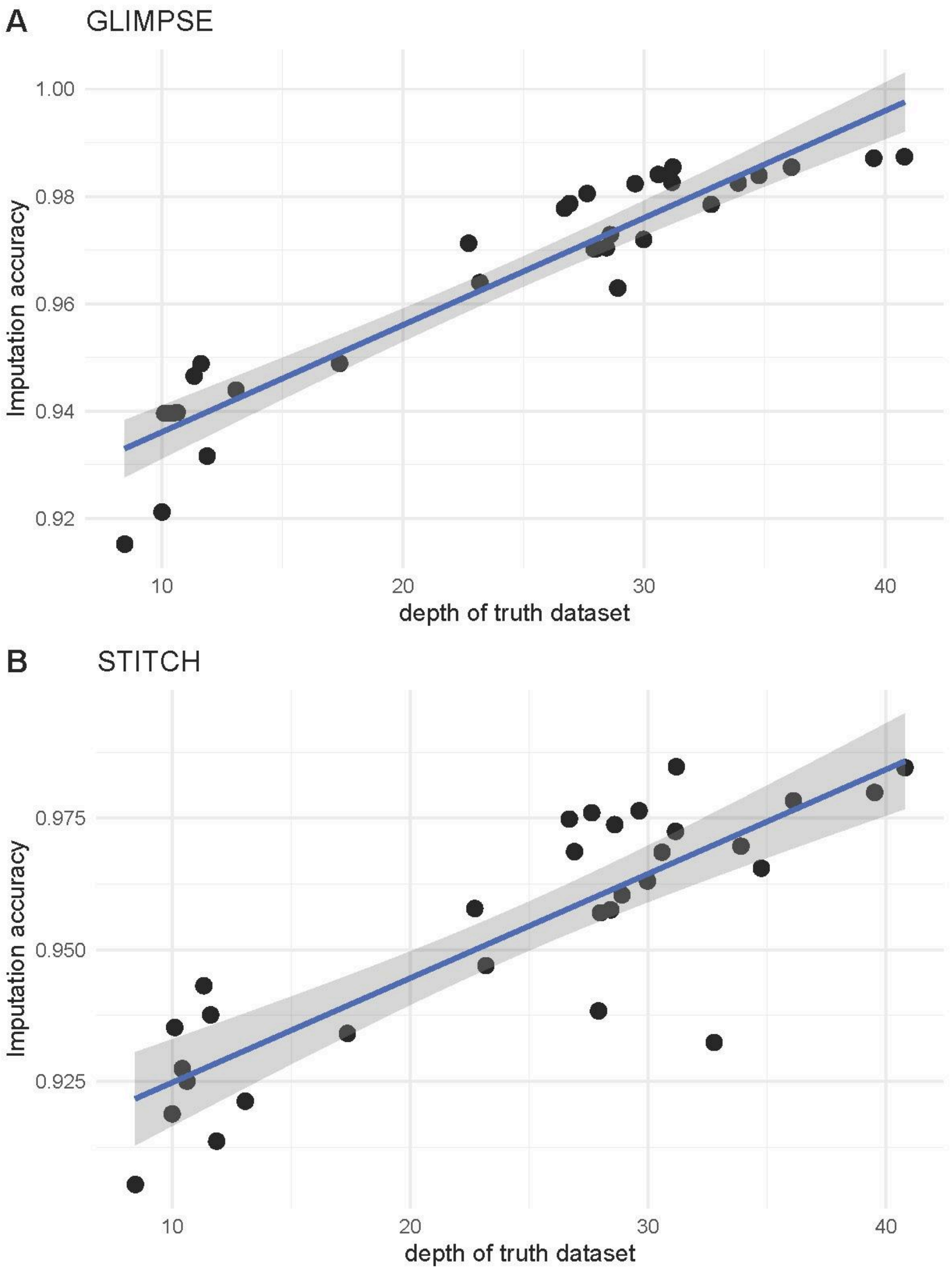

**Supplementary Figure 10 - Imputation accuracy as a function of the 'truth' sequencing depth.**

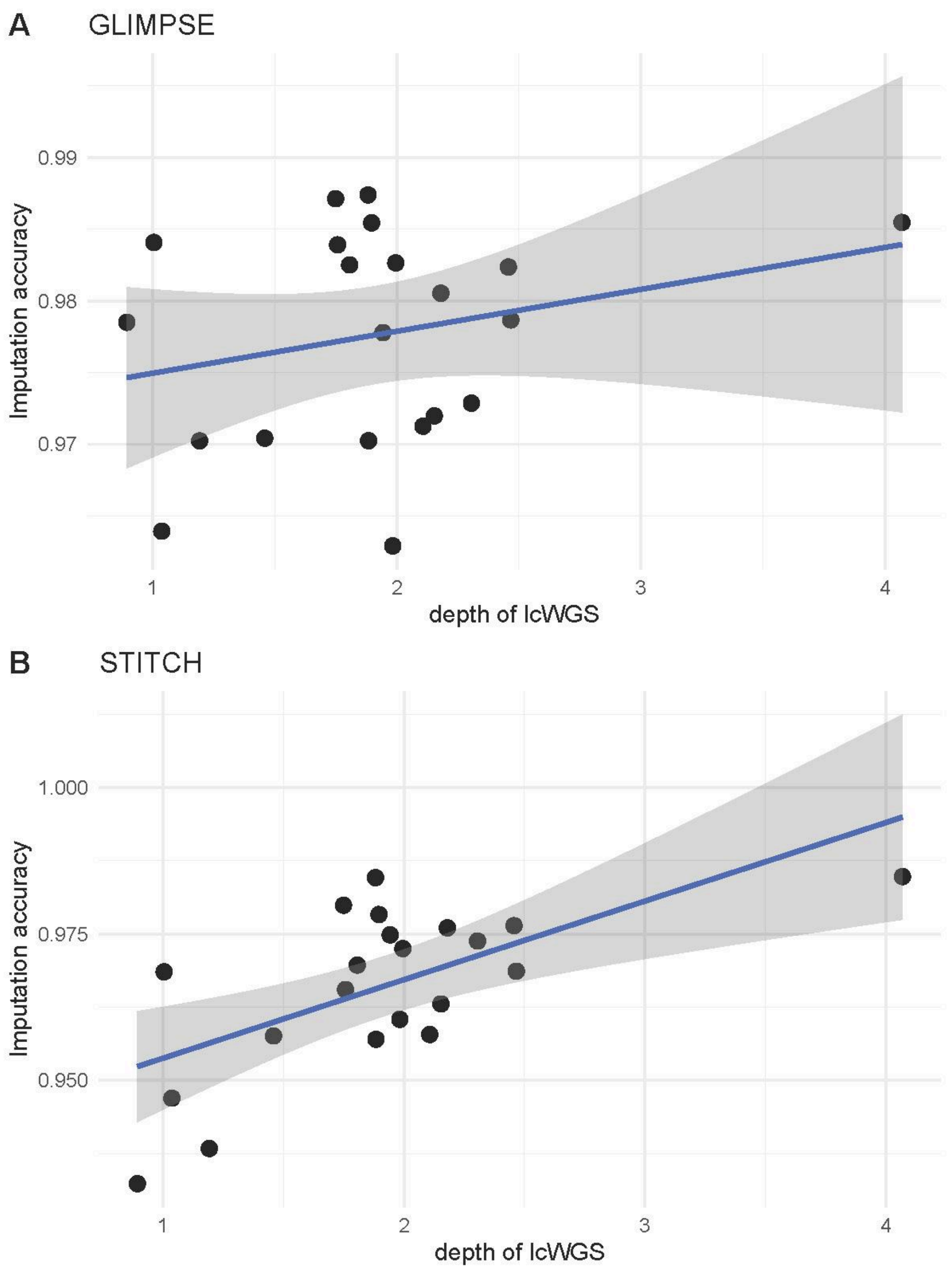

**Supplementary Figure 11 - Imputation accuracy as a function of the lcWGS sequencing depth.**

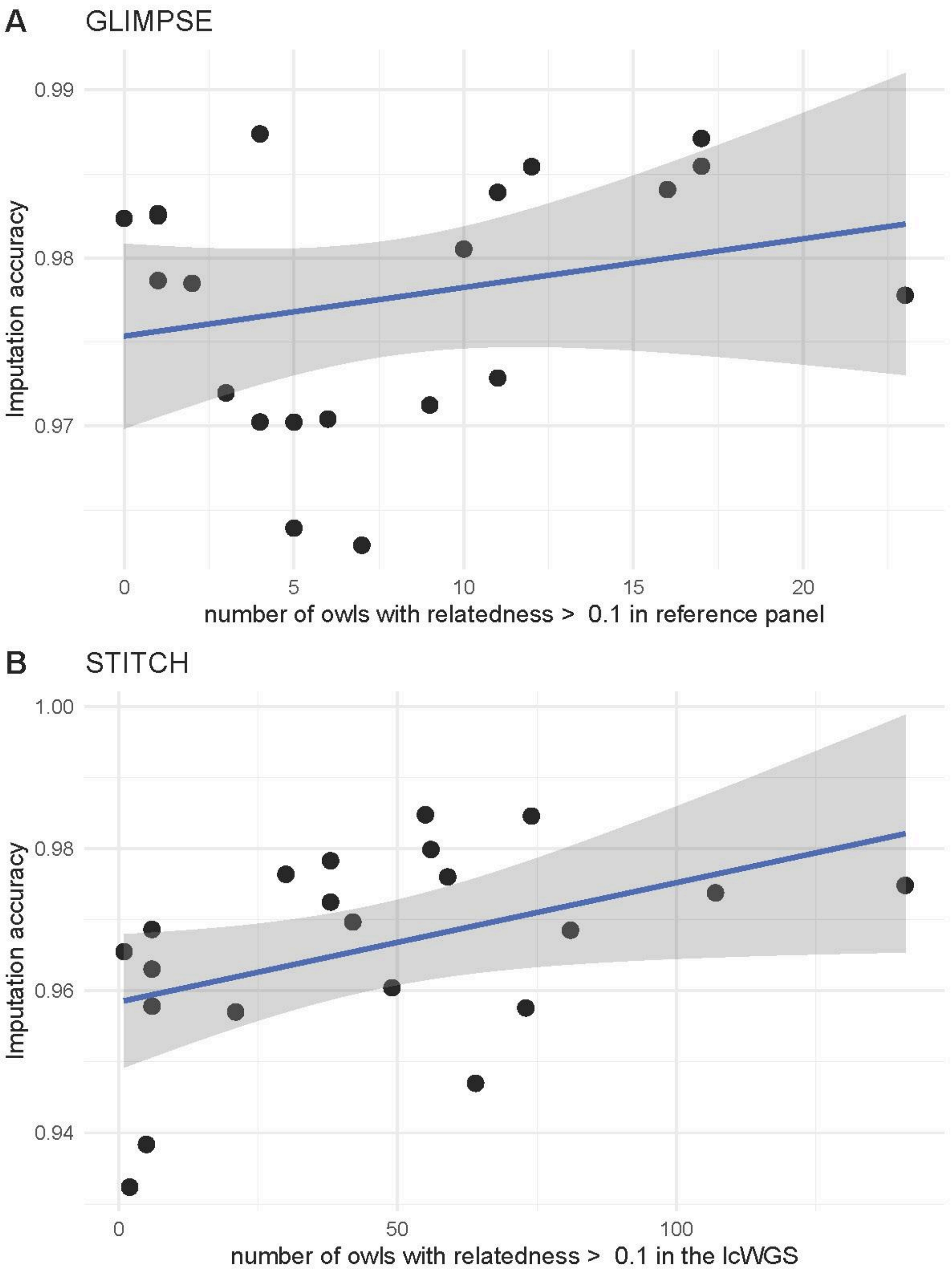

**Supplementary Figure 12. Imputation accuracy as a function of close relatives.**
**A)**. Imputation accuracy of GLIMPSE as a function of the number of close relatives (relatedness >0.1) between the 21 replicate individuals and the reference panel **B)** Imputation accuracy of STITCH as a function of the number of close relatives (relatedness > 0.1) between the 21 replicates and the lcWGS dataset

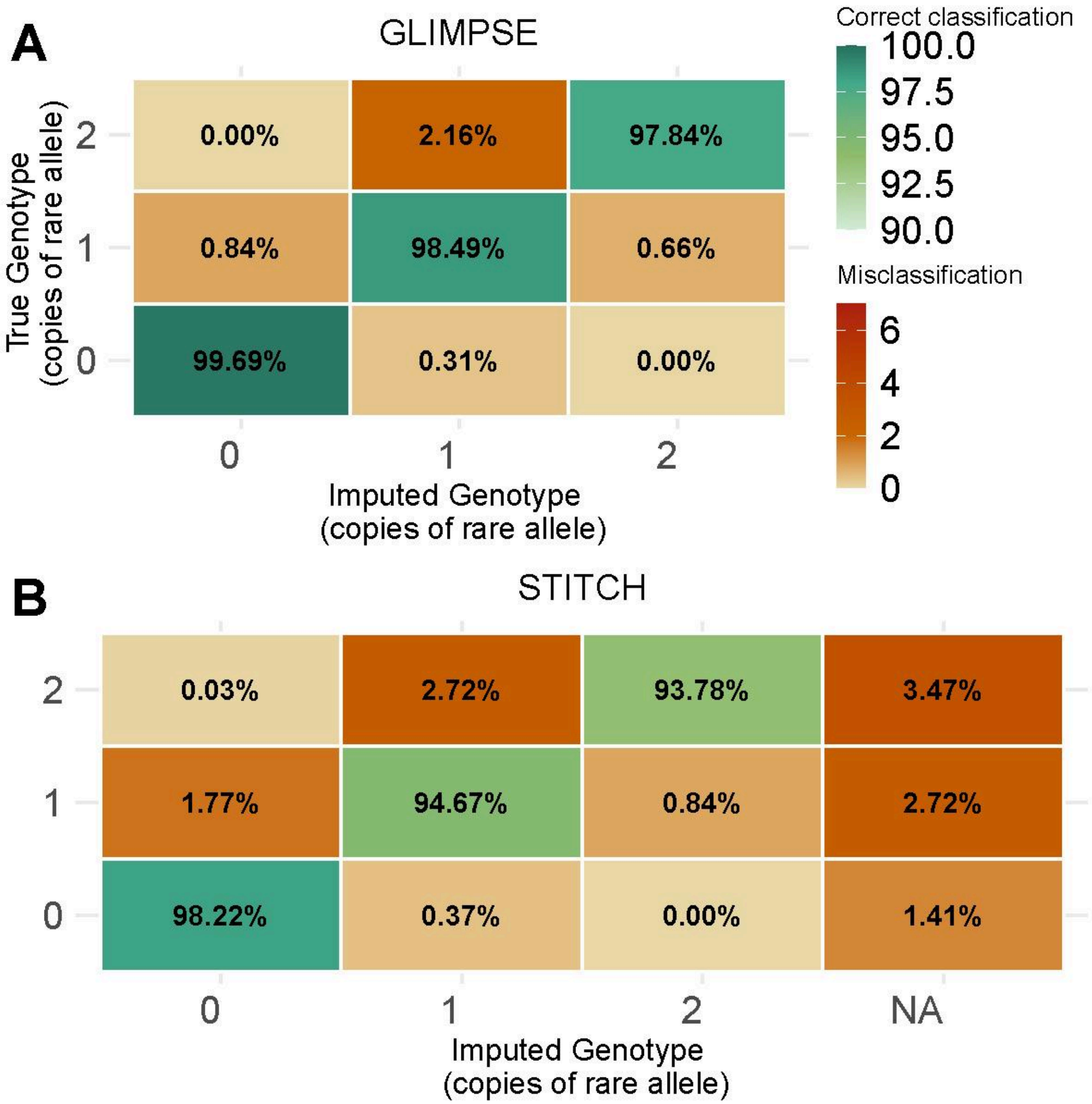

**Supplementary Figure 13. Misclassification profile across all allele frequencies for the two full datasets.** For GLIMSPE Each row represents the true genotype for these markers and the column the imputed one. A)For GLIMSPE B) Confusion matrix for genotype classification for STITCH. In B the NA column represents uncalled genotypes in STITCH. Diagonals are colored differently since higher is better, than off-diagonal where lower is better. All statistics calculated along the whole autosomal genome

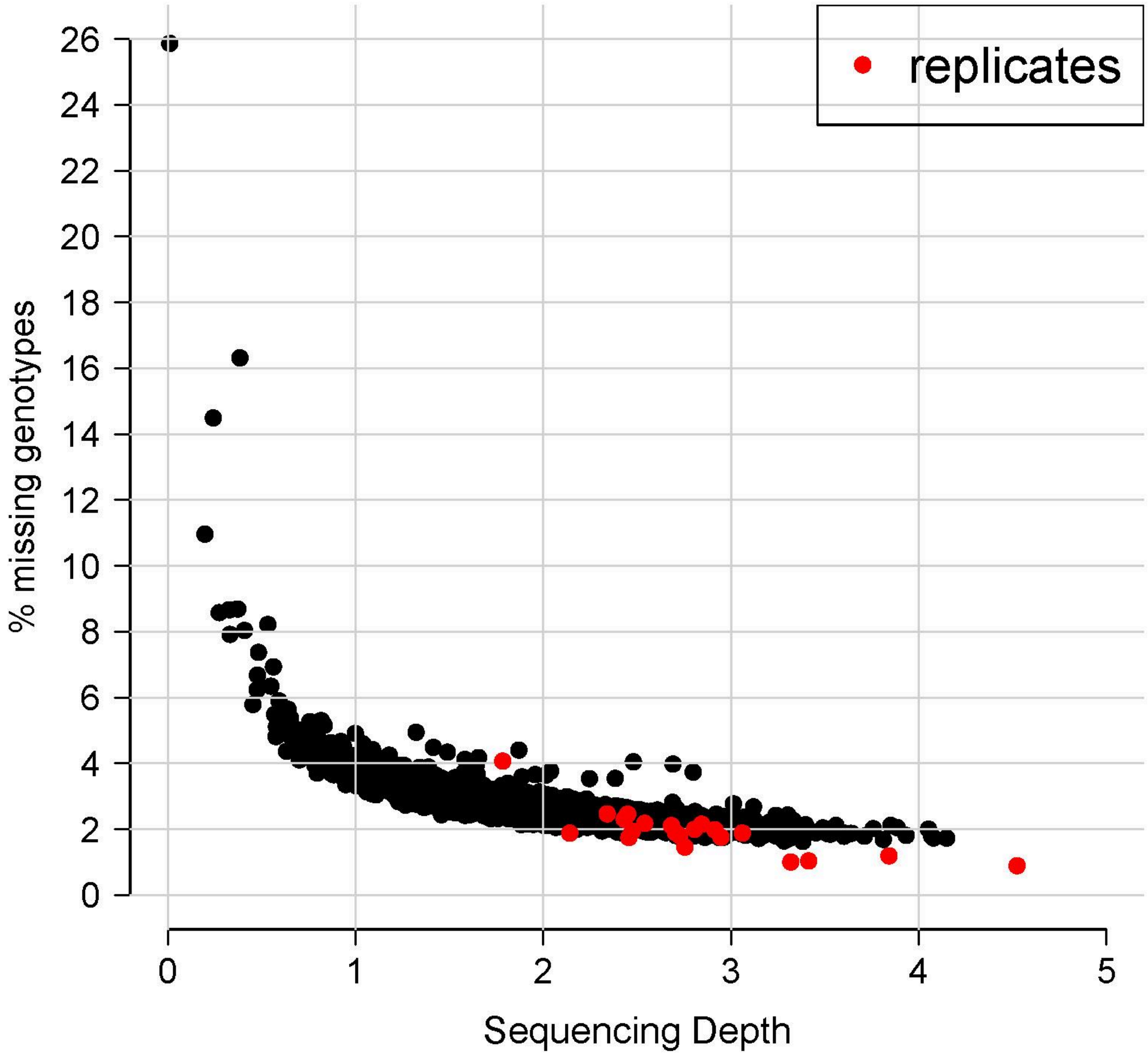

**Supplementary Figure 14 - Missing rate per sample in STITCH as a function of the sequencing depth.**
The n=21 replicates are highlighted with red.

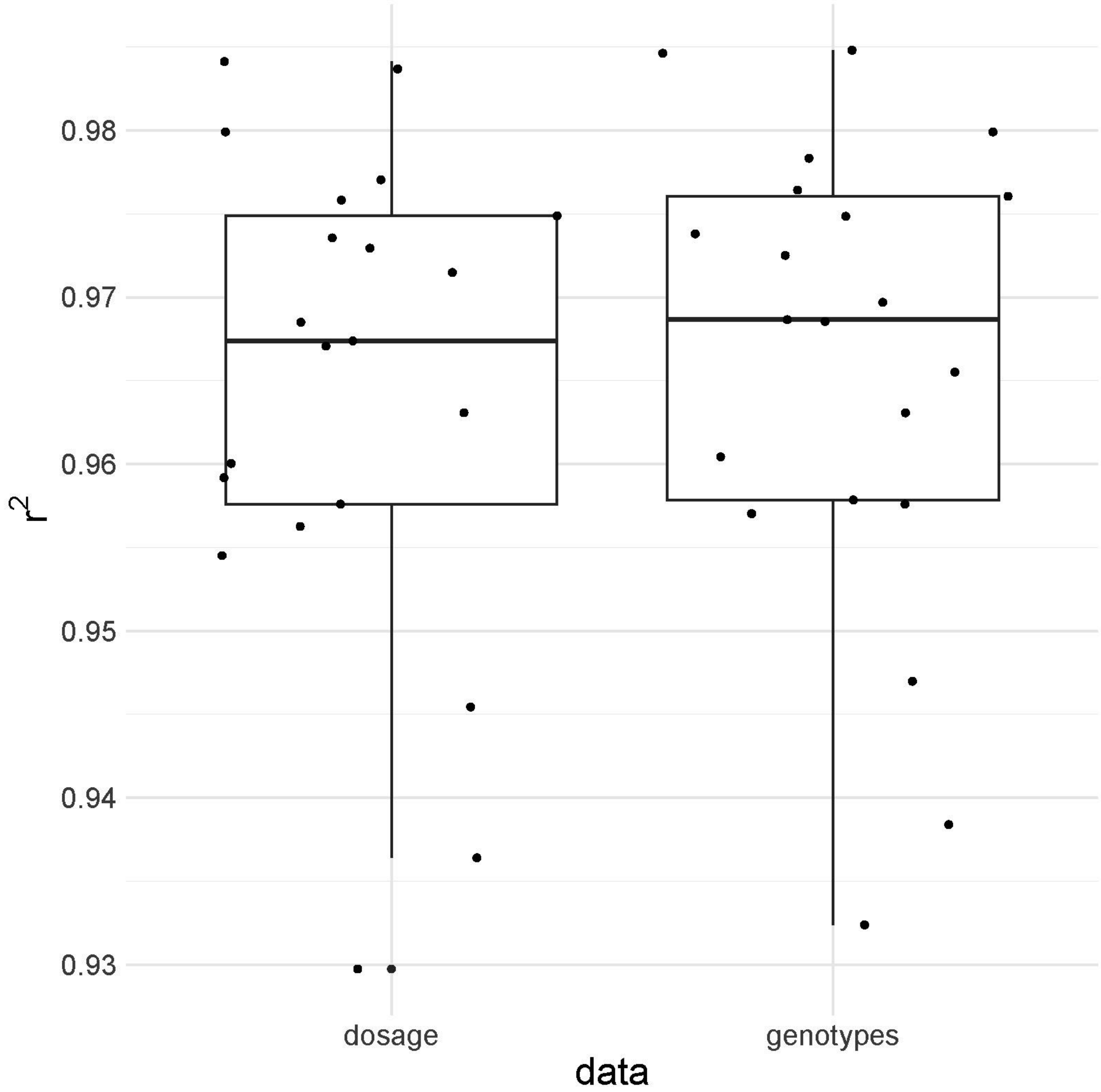

**Supplementary Figure 15 - Imputation accuracy with STITCH when using or not the missing genotypes.**
In ‘dosage’, we use the genotype dosage reported by STITCH which includes the genotypes set to missing by the software. On the contrary ‘genotypes’ use only the genotypes so it masks the missing data.

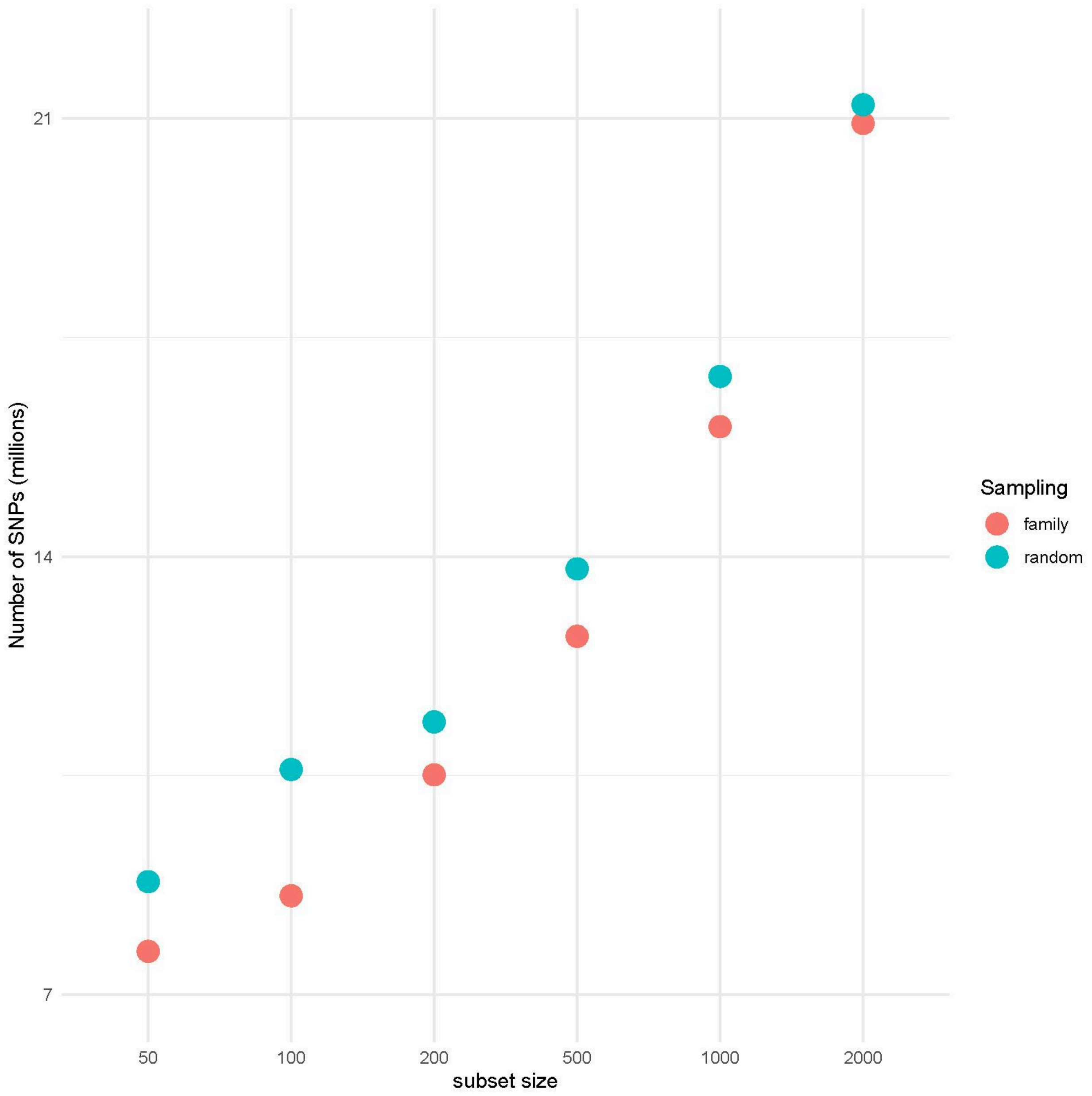

**Supplementary Figure 16 - Number of bi-allelic SNPs discovered in each subset.**

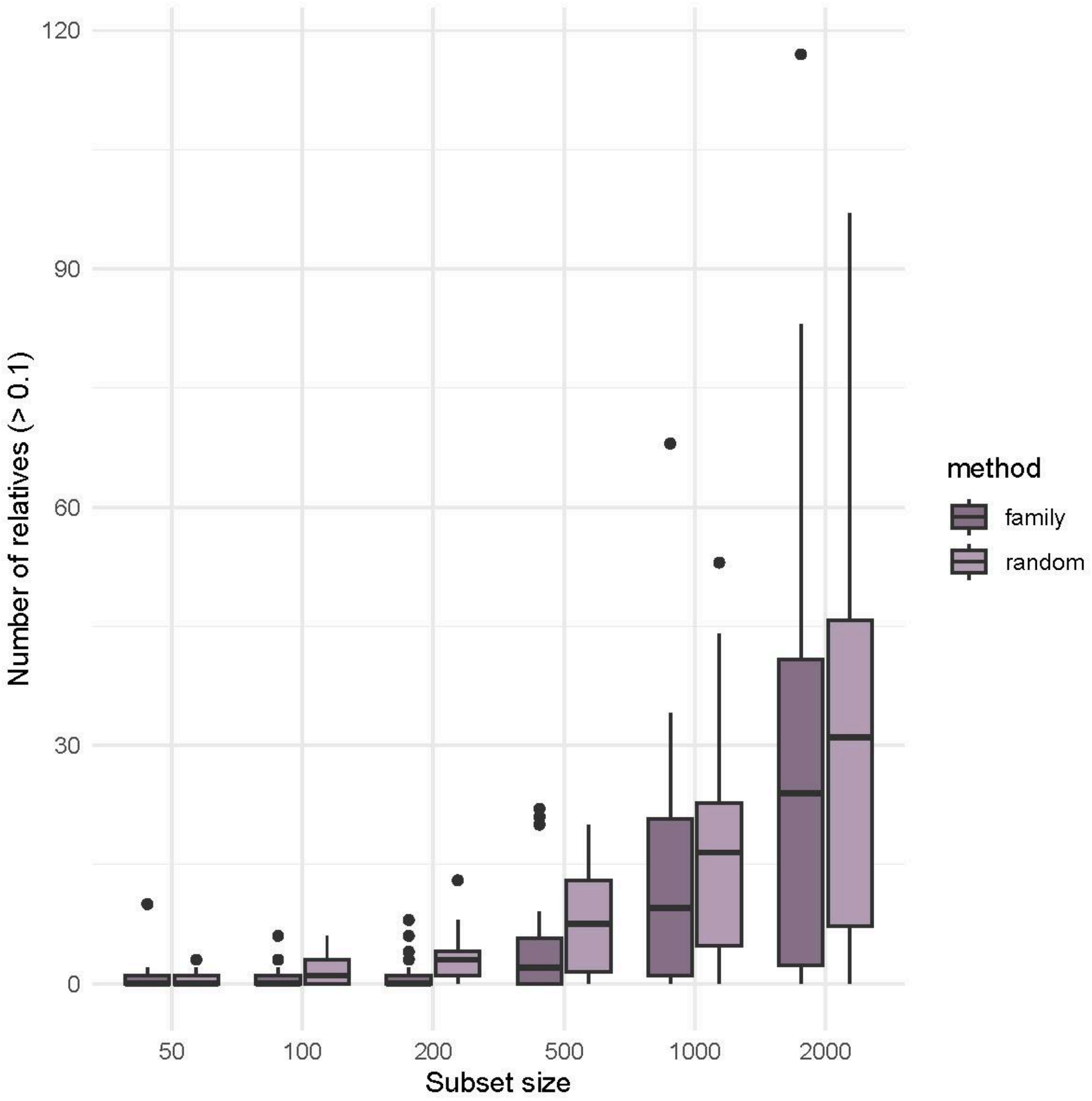

**Supplementary Figure 17**. The Number of 'close' relatives (relatedness >0.1) in each lcWGS subset dataset. l

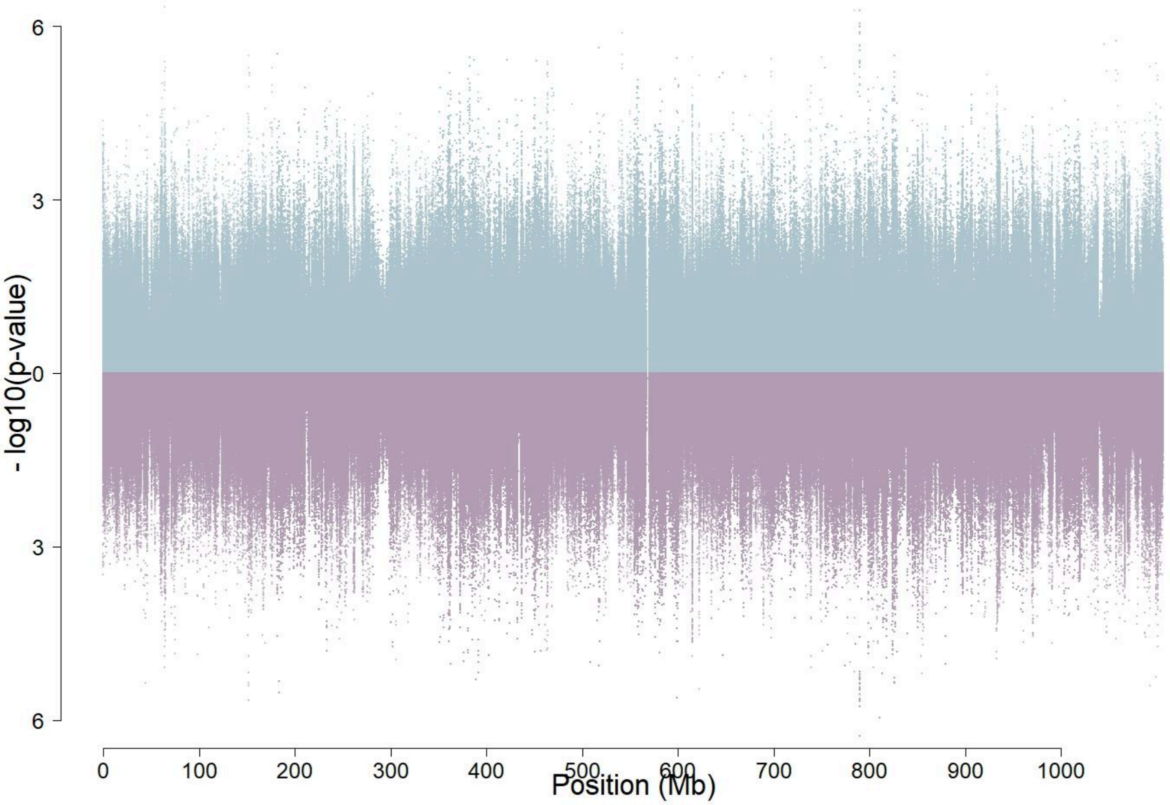

**Supplementary Figure 18.**
The whole genome GWAS on the size of the left tarsus of adult birds N=1980. Dots above the x-axis represent estimates with GLIMPSE. Dots below the x-axis represent estimates with STITCH.

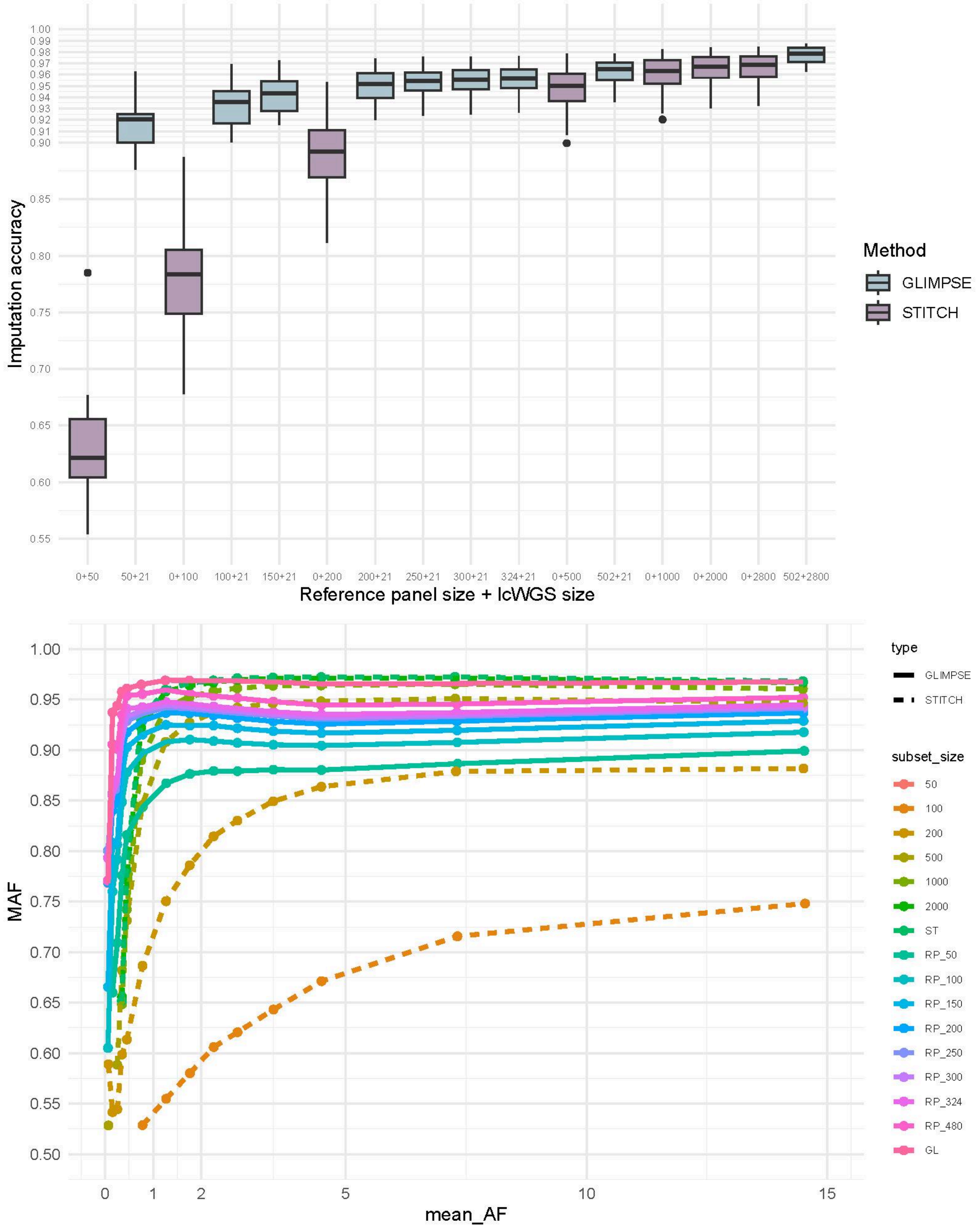

**Supplementary Figure 19**. The performance of all subseted and full datasets A) in individual imputation accuracy, B) in non-reference concordance across different allele frequencies.

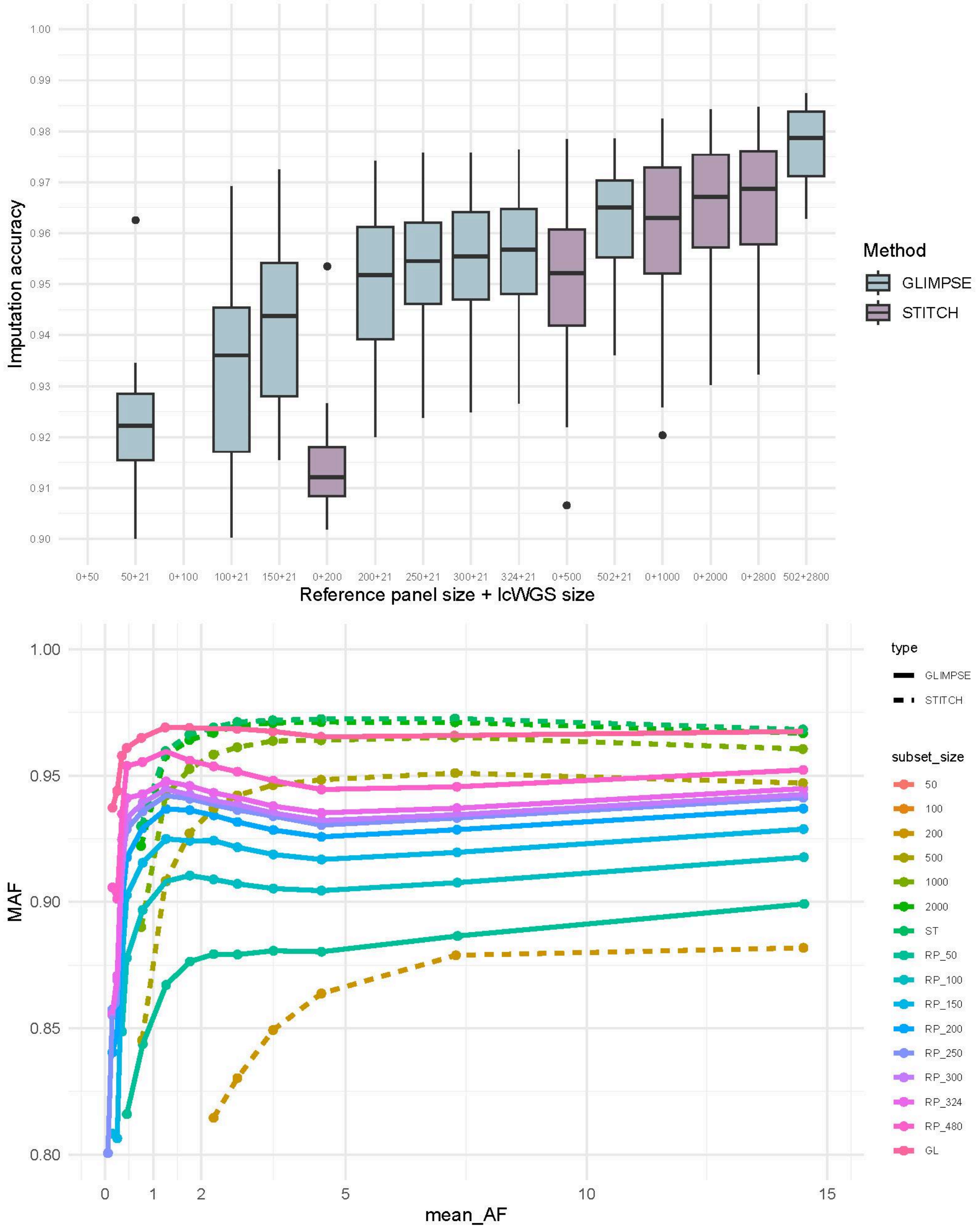

**Supplementary Figure 20**. The above figure with y-axis truncated for clarity. The performance of all subsets and full datasets A) in individual imputation accuracy, B) in non-reference concordance across different allele frequencies. In both panels, y-axis is truncated for visibility so some values may not appear for smaller datasets.

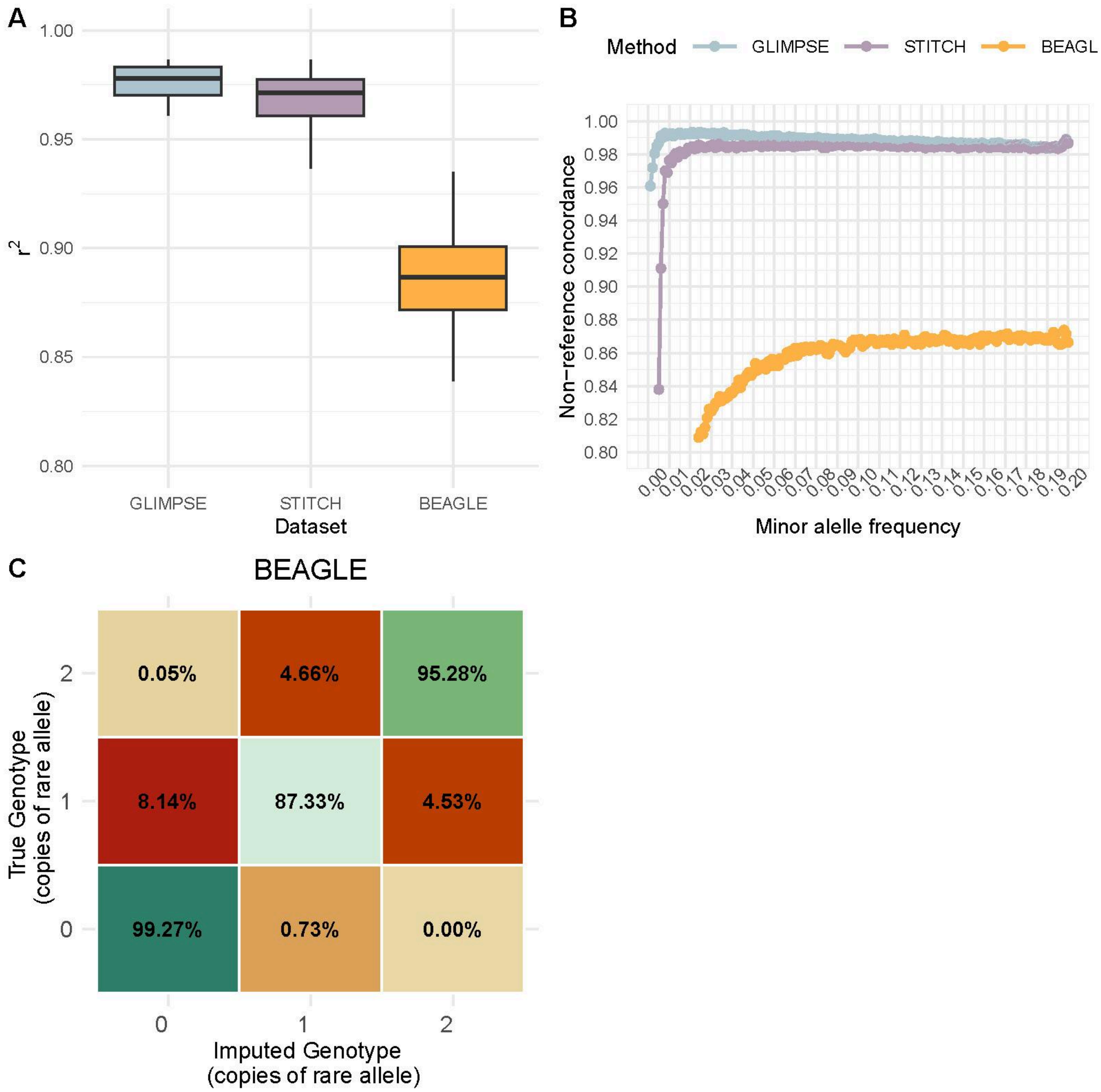

**Supplementary Figure 21. The imputation accuracy including the BEAGLE dataset**. A) Per sample imputation accuracy. B) The non-reference concordance (NRC) along different low-frequency alleles (effects estimated per SNP instead of in bins as in main text). C) The overall misclassification matrix when using BEAGLE.
